# SpatialFuser: a unified framework for integrative analysis of unpaired spatial multi-omics data

**DOI:** 10.1101/2025.09.14.676067

**Authors:** Wenhao Cai, Weizhong Li

## Abstract

Recent advances in spatial multi-omics technologies provide unprecedented opportunities to interpret molecular features in tissue microenvironments, but integrative analysis across heterogeneous datasets remains challenging. Here we present SpatialFuser, a deep learning framework for integrative analysis of unpaired spatial multi-omics data across epigenomics, transcriptomics, proteomics, and metabolomics. SpatialFuser consists of three coordinated modules: MCGATE, a Multi-head Collaborative Graph Attention auToEncoder that learns multi-scale spatial representations to decipher fine-grained spatial heterogeneity beyond predefined spatial neighbourhoods; an optional geometric pre-matching module that provides coarse initialization under tissue geometry mismatch; and an iterative matching–fusion module that couples geometry-constrained optimal transport matching with contrastive-learning-guided modality fusion for cross-slice alignment and integration. Systematic benchmarks demonstrate superior performance and reliability compared with existing state-of-the-art methods in spatial domain identification, cross-slice alignment, and multi-omics integration. Applications to real datasets illustrate that SpatialFuser resolves precise spatial molecular patterns, reveals developmental dynamics, and recovers complementary signals across modalities. Cross-resolution integration of weakly correlated modalities by our method further uncovers previously obscured biological variation. The generalizability and versatility of our framework enable customized analytical scenarios and potential extension for emerging omics.

**Highlights:**

- A unified deep learning framework for spatial multi-omics integrative data analysis
- Superior performance against state-of-the-art methods in spatial identification, alignment, and multi-omics integration
- Unprecedented cross-modality analysis scenarios to offer a holistic view of spatial multi-omics
- Comprehensive framework design with generalizability and versatility for customized scenarios and potential extension

## Introduction

Spatial technologies enable researchers to measure expression levels and reveal cellular heterogeneity across multiple molecular layers such as transcriptomics, epigenomics, proteomics, and metabolomics at specific spatial locations, and have been widely applied in studies of tissue organization and molecular dynamics[1]. These technologies can be roughly categorized into imaging-based approaches including osmFISH[2], MERFISH[3], CODEX[4], MIBI[5], CosMx SMI[6], and MALDI-MSI[7], and sequencing-based techniques such as 10X Visium[8], BaristaSeq[9], Stereo-seq[10], MAGIC-seq[11], Stereo-CITE-seq[12], Spatial ATAC-seq[13], and Spatial ATAC–RNA-seq[14]. Recent advances also cover AI-powered techniques such as PLATO[15], which leverage computational approaches to overcome current limitations in biotechnology, such as low sequencing depth. With different data modalities conveying unique biological insights from diverse complementary perspectives, we now have an unprecedented opportunity to achieve a comprehensive spatial landscape of cellular profile, thereby enhancing our understanding of key processes such as cell function[16], tissue development[17], and disease progression[18].

Computational methods have been developed to decipher intra-slice heterogeneity from typically spatial transcriptomics data with high-dimension and high-sparsity through spatial domain detection. SpaGCN[19] employs a graph convolutional network to integrate spatial gene expression with coordinate information for spatial domain identification. STAGATE[20] adopts a spatially-aware graph attention autoencoder to capture spatial structures, while SEDR[21] uses a deep variational autoencoder to jointly learn gene expression and spatial relationships. These unimodal methods model spatial relationships through predefined neighbourhoods, implicitly assuming that spatial proximity reflects biological similarity, an assumption that often breaks down in contexts such as tumour microenvironments, immune infiltration, and developmental migration. Meanwhile, the growing expansion of spatial technologies into multi-omics indicates that increasing numbers of tissue slices will be profiled from proteomic, epigenomic, and other molecular perspectives, underlying this is an urgent need for unified computational tools capable of decoding spatial data from diverse omics modalities.

To achieve a deeper understanding of complex biological systems, integrative analysis of cross-modality spatial data has become increasingly important. However, heterogeneous datasets often differ in tissue geometry, spatial resolution and cellular composition, while batch effects and modality-specific biases further introduce discrepancies in feature structures, expression distributions and biological semantics, posing challenges for effective data integration and slice alignment. Existing computational tools attempt to address these issues but remain limited. For example, SIMO[22] employs probabilistic alignment to map single-cell multi-omics data onto homologous spatial transcriptomics slices for detecting spatial cellular topological patterns, but it requires paired datasets and is restricted to within-slice analysis. STAligner[23] employs a graph attention autoencoder and triplet adversarial learning to align spatial transcriptomic slices, but it does not extend to multi-omics integration. SLAT[24] uses graph convolutional networks and adversarial learning to reconstruct 3D tissues while correcting batch effects across slices, yet it fails to integrate modalities with weakly correlated features, such as transcriptomics and proteomics.

In the presence of substantial feature semantic discrepancies across modalities, explicitly modeling feature dependencies becomes challenging. MISO[25] addresses this by learning modality-specific embeddings and their interactions to identify biologically relevant spatial domains, but it cannot perform cross-slice integration. Similarly, SpatialGlue[26] and MultiGATE[27] apply dual-attention graph models to jointly analyse spatial multi-omics from the same tissue, yet they heavily rely on shared spatial coordinate systems or hard-coded biological priors to link weakly correlated features. These integrative methods are effective for unimodal or same-slice multi-omics data but lack generalizability in unpaired multi-omics scenarios, where datasets do not share matched spatial coordinates and often differ in modality, resolution, geometry, or cellular composition. Methodologically, many integrative tools assume consistent feature structures across datasets, requiring dimensionality matching through preprocessing. This can lead to information loss when the intrinsic complexity of raw modalities differs. Some also assume similar cell compositions across slices and adopt rigid integration strategies, increasing the risk of over-alignment. Therefore, new computational tools are urgently needed for spatially aware cross-omics integration that can offer better biological interpretability and generalizability through more flexible and robust analysis across diverse and emerging spatial omics modalities.

Here, we introduce SpatialFuser, a unified deep learning framework for integrative analysis of unpaired spatial multi-omics data, enabling accurate spatial interpretation, effective cross-slice alignment and robust cross-modality integration. Rather than relying solely on fixed spatial neighbourhood structures, SpatialFuser learns multi-scale cellular associations through a collaborative graph attention mechanism. This design preserves local spatial continuity while incorporating weighted long-range links between molecularly similar but spatially distant spots, thereby complementing local neighbourhood aggregation for more informative spatial representation learning. For cross-slice analysis, SpatialFuser optionally incorporates geometric pre-matching for coarse initialization under geometric mismatch. Rather than treating alignment and integration as separate sequential procedures, SpatialFuser formulates them as a coupled optimisation problem. By iteratively updating geometry-constrained optimal transport matching and contrastive fusion, SpatialFuser jointly refines cross-slice correspondences and shared representations across unpaired slices with different modalities, resolutions, and tissue geometries.

We benchmarked SpatialFuser on classical 10X Visium datasets to evaluate its quantitative accuracy and robustness in spatial domain identification, and then validated its cross-platform and cross-modality generalizability on osmFISH and CODEX datasets. In unimodal multi-slice joint analysis, SpatialFuser achieved better alignment and integration performance than state-of-the-art methods reported in previous benchmarks[28] on adjacent BaristaSeq slices, and effectively captured fine-grained developmental dynamics through integrative analysis of multi-developmental-stage spatial Stereo-seq samples. In spatial multi-omics scenarios, SpatialFuser outperforms existing methods in resolving subtle spatial patterns of cellular states and enhances the detection of lowly expressed but epigenetically primed marker genes by integrating complementary information from spatial ATAC–RNA-seq data, thereby revealing lineage-associated regulatory signals. In integrative analysis across multi-omics samples with weakly correlated features, SpatialFuser effectively integrates transcriptomic (MAGIC-seq), proteomic (PLATO), and metabolomic (MALDI-MSI) data across varying spatial resolutions and mismatched tissue geometries to improve the fine-grained characterization of spatial topological patterns. These results highlight the capability of SpatialFuser to infer cell state transitions and transcriptional readiness within spatial tissue architectures, and to identify fine-grained functional regions that are challenging to resolve with low-resolution technologies alone. SpatialFuser demonstrates versatility and generalizability in decoding complex biological systems through comprehensive spatial multi-omics integration.

## Results

### The overview architecture of SpatialFuser

SpatialFuser is a deep-learning framework designed for both fine-grained single-slice analysis and cross-sample integrative analysis across spatial epigenomic, transcriptomic, proteomic, and metabolomic data (Fig. 1a), supporting tasks including spatial interpretation, cross-slice alignment, and cross-modality integration. It takes molecular features and spatial coordinates as input, and first represents each spatial omics slice as graph-structured data by constructing a spatial adjacency network from the coordinates. To accommodate multi-omics datasets with different feature dimensions and molecular semantics, SpatialFuser employs a Multi-head Collaborative Graph ATtention autoEncoder (MCGATE) to learn slice-specific embeddings through a reconstruction task, rather than enforcing a shared encoder across heterogeneous inputs (Fig. 1b Step 1).

**Fig. 1.**
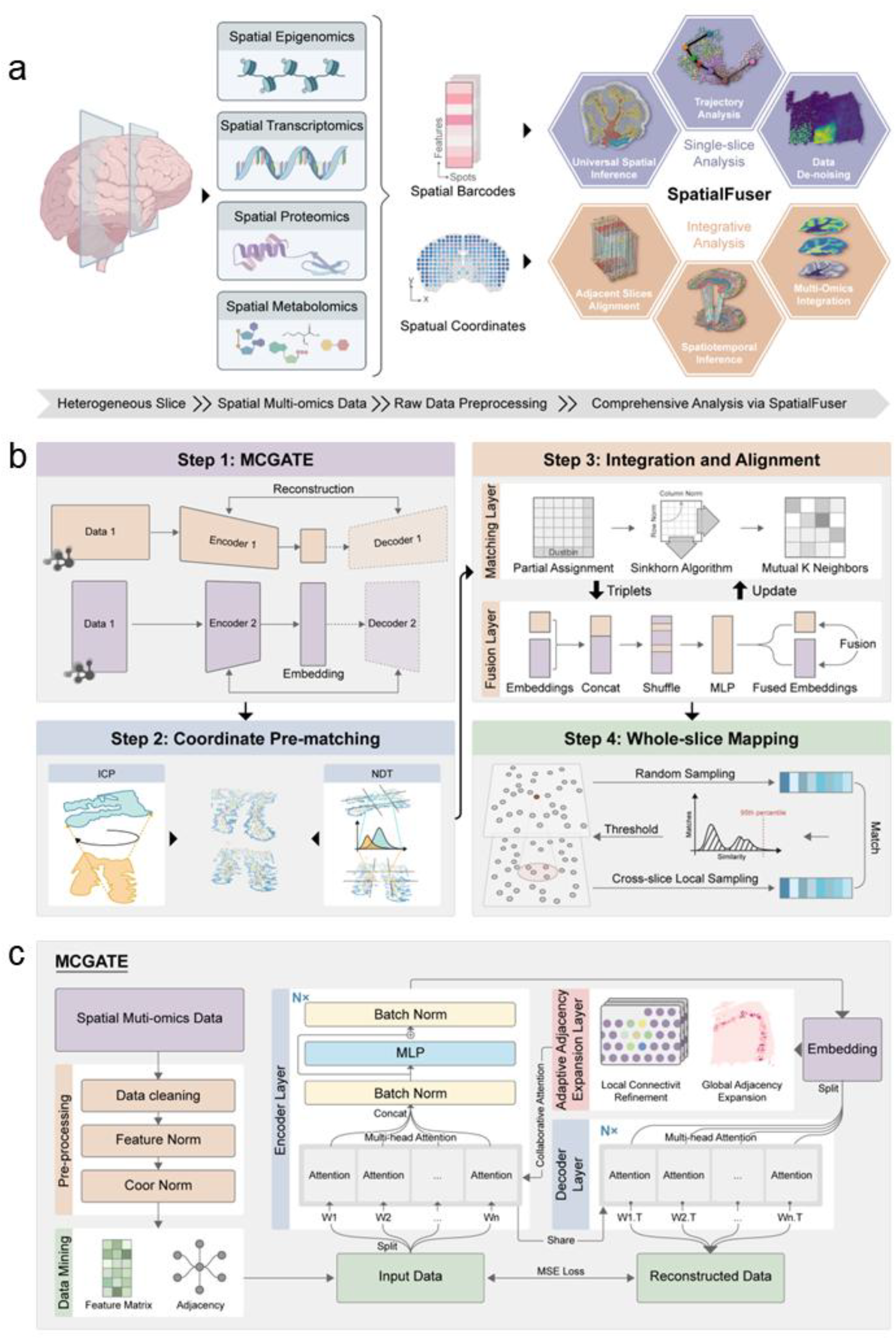
A unified deep learning framework for single-sample and cross-slice spatial multi-omics analysis. a, Overview of SpatialFuser. SpatialFuser integrates molecular features and spatial coordinates from spatial omics data, supporting both fine-grained single-slice profiling and cross-sample analysis across modalities such as epigenomics, transcriptomics, proteomics, and metabolomics. b, Schematic representation of the SpatialFuser framework. The analysis workflow includes: MCGATE-based embedding learning, coordinate pre-matching for rigid slice registration, dual matching and fusion layers for multimodal integration, and whole-slice spot mapping for downstream integrative analysis. c, Network architecture of MCGATE. MCGATE represents spatial omics data as a graph by constructing a spatial adjacency network from coordinates. It applies multi-head collaborative graph attention to learn modality-specific embeddings via a reconstruction task, guided by a spatial graph for local attention and a dynamically updated feature similarity graph for long-range attention. Designed to be modular and scalable, MCGATE can be applied across diverse spatial omics modalities and platforms.

In MCGATE (Fig. 1c), the predefined spatial adjacency graph serves as the local attention aggregation backbone, while a latent feature similarity graph introduces controlled long-range attention links between molecularly similar but spatially distant spots. This collaborative attention design enables MCGATE to model spatial context at multiple scales and learn more informative representations of tissue organization. As a general embedding model for spatial omics data, MCGATE remains modular and scalable, supporting its application across diverse modalities and experimental platforms.

When substantial geometric mismatch exists across tissue slices, coordinate pre-matching can be critical for providing a coarse geometric initialization for downstream spatial omics alignment and integration[29]. To accommodate diverse spatial distributions, SpatialFuser models spatial omics data as 2D point clouds and provides an optional coordinate registration module based on Iterative Closest Point (ICP) and Normal Distributions Transform (NDT) algorithms (Fig. 1b Step 2).

Alignment and integration are not strictly independent tasks[28]. SpatialFuser therefore couples them through an iteratively trained dual-layer architecture consisting of an optimal transport-based matching layer and a contrastive-learning-based fusion layer. (Fig. 1b Step 3). In the matching layer, cross-slice alignment is formulated as a graph matching task and solved through Sinkhorn-based optimal transport. Spatial geometry constrains the candidate matching space, providing a structural prior when cross-slice representations are not yet fully aligned during early training. Within this constrained space, the Sinkhorn algorithm generates soft cross-slice matching scores, from which high-confidence matched pairs are selected for downstream integration. To avoid forcible alignment under cell-type compositional or resolution mismatch, a dustbin channel mechanism[30] is incorporated to filter out ambiguous or low-confidence assignments. In the fusion layer, independent MCGATE embeddings are first projected into a shared latent space. High-confidence matched pairs from the matching layer are used as anchor-positive pairs for contrastive learning, while anchor-negative pairs are sampled from non-neighbouring regions within the same slice. This contrastive objective aligns matched cross-slice pairs while separating within-slice negatives, reducing batch effects and modality biases without collapsing biological variation. The updated shared latent space is subsequently used to refine matching in the next iteration, allowing alignment and modality fusion to improve jointly. Finally, SpatialFuser applies quality control to achieve whole-slice spot mapping (Fig. 1b Step 4), providing a reliable reference for downstream integrative analyses.

In brief, SpatialFuser comprises three main modules: MCGATE, optional coordinate pre-matching, and an iterative matching–fusion module consisting of an optimal transport-based matching layer and a contrastive fusion layer. In this framework, MCGATE provides a general embedding engine for single-slice spatial interpretation across diverse platforms and modalities. Building on these slice-specific representations, the matching–fusion module further couples cross-slice alignment with modality fusion, enabling integrative modelling of unpaired and heterogeneous spatial multi-omics datasets.

### Accurate spatial inference across diverse experimental platforms and modalities

Accurate single-slice spatial interpretation is a prerequisite for reliable multi-slice and multi-omics integrative analysis. We therefore first examined the single-slice representation capacity of SpatialFuser across diverse platforms and modalities. To evaluate the performance of SpatialFuser against existing representation learning algorithms, we conducted a benchmark on spatial domain detection task using the classic 10X Visium[8] human dorsolateral prefrontal cortex (DLPFC) dataset[32], which has well-defined morphological boundaries and reliable manual annotations (Fig. 2a & Supplementary Fig. 1). The original labels were used as ground truth, and five evaluation metrics were employed: Adjusted Rand Index (ARI), Adjusted Mutual Information (AMI), Homogeneity, Completeness, and V-Measure. We first compared SpatialFuser with other state-of-the-art spatial domain detection methods based on the Mclust[33] clustering strategy, including SpaGCN[19], STAGATE[20], SEDR[21], and STAligner[23] (Fig. 2b). On average across the 12 DLPFC slices, SpatialFuser achieved the highest accuracy (mean ARI = 0.625; max = 0.809; min = 0.533), followed by STAligner (mean ARI = 0.556; max = 0.675; min = 0.469). Notably, although SpaGCN integrates high-resolution histological images during training, its performance was substantially lower than that of other methods relying solely on molecular and spatial information (mean ARI = 0.422; max = 0.541; min = 0.225), suggesting that the utility of histological information is highly dependent on the modeling and integration strategy, rather than on its inclusion alone. To further evaluate SpatialFuser’s advantages in downstream analysis, we also benchmarked it using commonly used clustering methods in omics data analysis, including Leiden[34] (Supplementary Fig. 2a) and Louvain[35] (Supplementary Fig. 2b). These results demonstrate that SpatialFuser enables accurate spatial domain identification and consistently outperforms existing state-of-the-art tools across diverse clustering contexts.

**Fig. 2.**
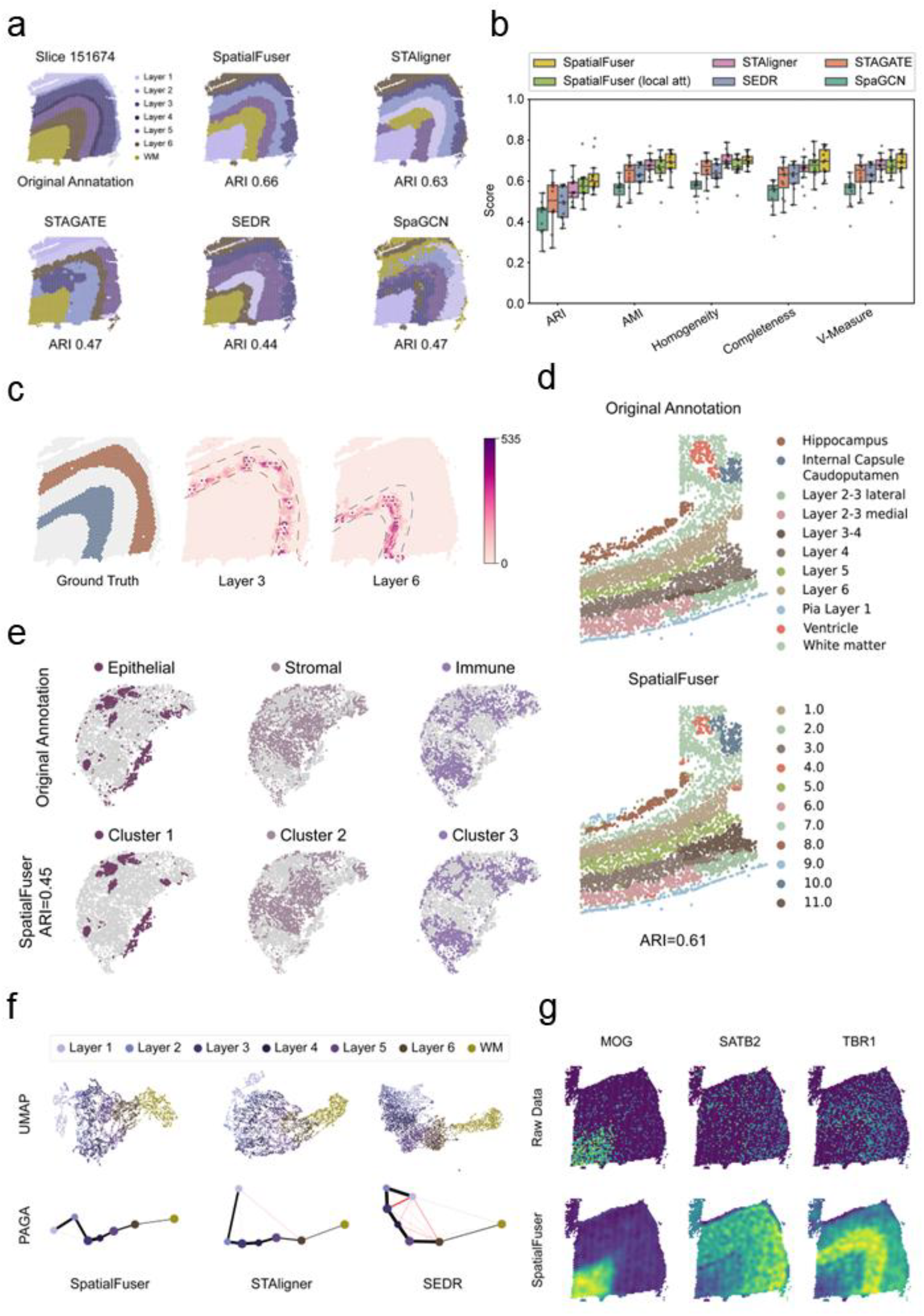
SpatialFuser accurately resolves tissue domains across diverse experimental platforms and modalities. **a**, Comparison of spatial domains identified by SpaGCN, STAGATE, SEDR, STAligner, and SpatialFuser for DLPFC slice-151674. **b**, Boxplots of five evaluation metrics (ARI, AMI, Homogeneity, Completeness, V-measure) for Mclust clustering results across 12 DLPFC sections. **c**, Long-range feature propagation paths of ten randomly selected spots from cortical layers 3 and 6 during MCGATE training on DLPFC slice-151676. **d**, Spatial domains identified by Mclust clustering on low-dimensional embeddings generated by SpatialFuser in the osmFISH somatosensory cortex dataset. **e**, Distributions of epithelial, stromal, and immune cells identified by Mclust clustering on low-dimensional embeddings generated by SpatialFuser for slice-210308_TMA2_reg6 of the CODEX muscle-invasive bladder cancer dataset. **f**, UMAP visualizations and PAGA graphs generated from SEDR, STAligner, and SpatialFuser embeddings for DLPFC section-151675. Red lines highlight incorrect trajectory inferences. **g**, SpatialFuser enhances spatial patterns of layer-enriched and marker genes in the DLPFC dataset. Raw and SpatialFuser-denoised spatial expression patterns are shown for MOG, SATB2, and TBR1 in DLPFC section-151675. ARI, adjusted Rand index; AMI, adjusted mutual information; DLPFC, dorsolateral prefrontal cortex; MCGATE, multi-head collaborative graph attention autoencoder.

We conducted an ablation study to evaluate the effectiveness of the collaborative attention mechanism. The results demonstrate that the long-range attention significantly enhances the SpatialFuser’s ability to capture spatial distribution patterns (Fig. 2b, Supplementary Fig. 2a-b). Using slice-151676 from the DLPFC dataset as an example, we randomly selected 10 spatial spots from cortical layers 3 and 6 and tracked their long-range correspondence sampling during training. The distribution of sampled spots closely aligned to the tissue region patterns (Fig. 2c), further validating the effectiveness and biological relevance of long-range information propagation. Additional attention-head ablation on the DLPFC benchmark showed improved clustering performance under multi-head configurations compared with the single-head setting (Supplementary Fig. 3). We also tested the robustness of SpatialFuser by comparing clustering accuracy across different hyperparameters (see Methods), demonstrating that the model is insensitive to network structure and random seeds (Supplementary Fig. 3a-b).

Spatial omics data generated from different experimental platforms and modalities exhibit significant differences in feature structures, distributions, and biological semantics. We further evaluated SpatialFuser’s ability to identify spatial domains across multiple platforms and modalities using the osmFISH somatosensory cortex dataset[2] and the CODEX[4] human muscle-invasive bladder cancer tumor dataset[36]. On the image-based osmFISH dataset, SpatialFuser accurately identified spatial domains in the somatosensory cortex with an ARI of 0.61 (Fig. 2d). For the CODEX spatial proteomics dataset, we re-labelled the three main cell types, epithelial cells, stromal cells, and immune cells, mentioned in the original paper based on the annotations provided by the authors for downstream spatial inference. Although tumour heterogeneity may reduce the agreement between cell-type labels and discrete clustering metrics such as ARI[37], SpatialFuser successfully captured the spatial distribution patterns of all three key cell types (Fig. 2e, ARI = 0.45). These results demonstrate the robustness and versatility of SpatialFuser in cross-platform, multi-omics spatial data analysis.

To evaluate SpatialFuser’s utility in downstream analysis, we assessed its low-dimensional embeddings for trajectory inference and reconstructed features for spatial variable gene (SVG) identification. Partition-based graph abstraction[38] (PAGA) was employed to infer trajectories on DLPFC slice-151675 using embeddings learned by SpatialFuser, SEDR, and STAligner. In the UMAP plot (Fig. 2f), SpatialFuser’s embedding closely recapitulated the developmental progression from white matter to layer 6 and sequentially to layer 1. Comparatively, embeddings from SEDR and STAligner exhibited less plausible developmental trajectories. We further compared the expression patterns of three region-enriched marker genes between the raw data and SpatialFuser-denoised data in DLPFC slice-151675. After reconstruction, all selected genes showed clear differential expression in their enriched regions, while the raw spatial expression was noisy and inconsistent (Fig. 2g). Collectively, these results demonstrate that SpatialFuser can accurately capture feature expression patterns and reduce noise, thereby better supporting single-sample analysis and spatial heterogeneity studies.

### Spatiotemporal alignment for revealing tissue heterogeneity and developmental dynamics

Integrative and comparative analysis of spatial omics datasets is critical for unravelling spatial complexity and temporal changes in tissue systems[39]. To assess the performance of SpatialFuser in pairwise slice alignment and integration, we first applied it to two adjacent slices from the BaristaSeq mouse visual cortex dataset[9], comprising 1525 and 2042 spots, respectively (Fig. 3a). We evaluated spot-to-spot alignment accuracy across the entire tissue section (see Methods), comparing SpatialFuser with three state-of-the-art methods reported in previous benchmarks[28], including SPACEL[40], SLAT[24], and STAligner[23] (Fig. 3b). Owing to its use of ground-truth labels during training (see Methods), SPACEL achieved the highest alignment accuracy (0.987, 1519 matches), followed closely by SpatialFuser (0.972, 1516 matches) and STAligner (0.949, 1508 matches), while SLAT showed noticeably lower performance (0.857, 1525 matches).

**Fig. 3.**
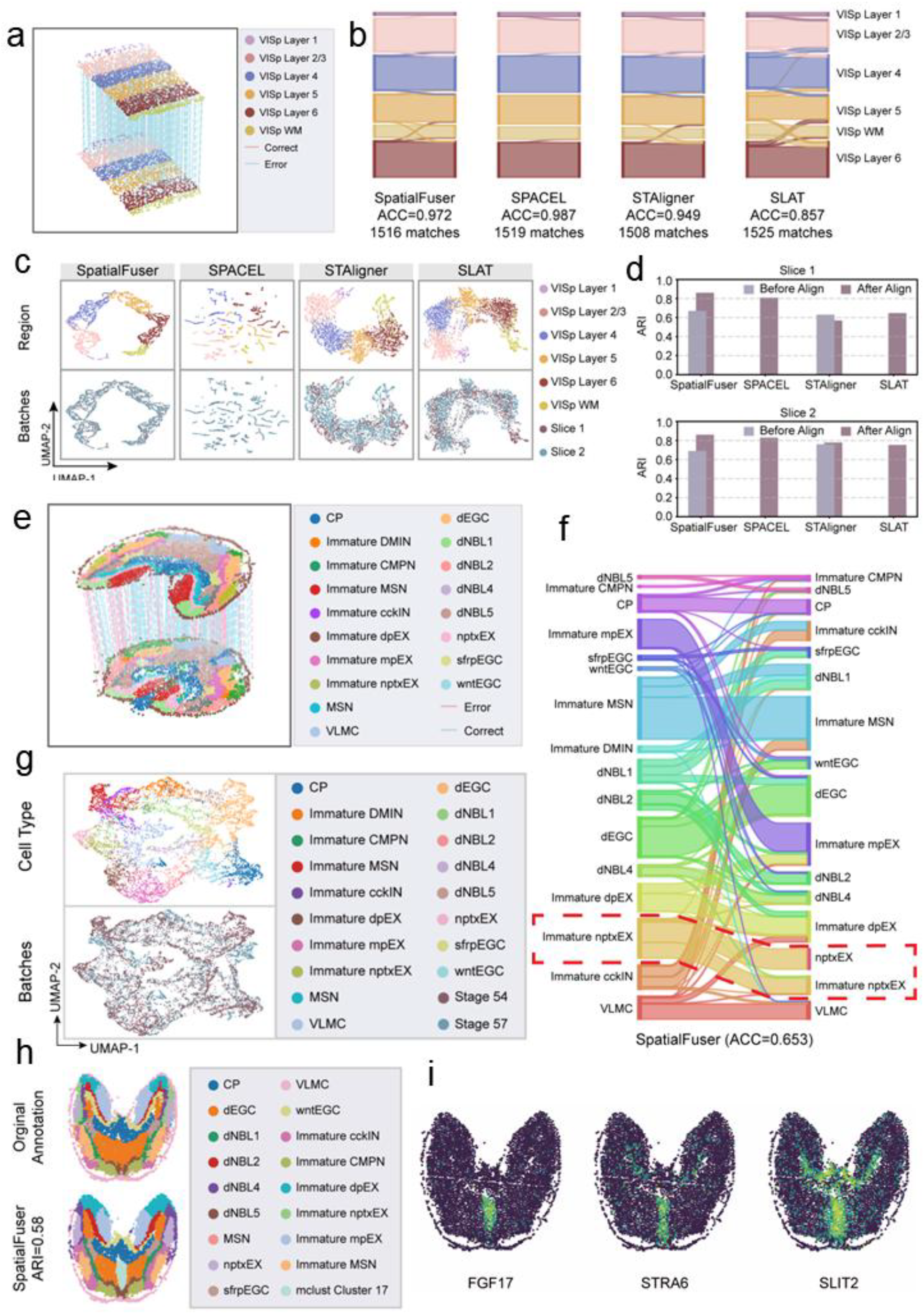
Spatial alignment across consecutive tissue slices and non-consecutive slices from different developmental stages. **a**, Alignment of slices 1 and 2 from the BaristaSeq mouse visual cortex dataset, coloured by region labels (300 alignment pairs shown for clarity). **b**, Sankey plot showing region type correspondence based on alignments from SpatialFuser, SPACEL, STAligner, and SLAT for slices 1 and 2 of the BaristaSeq dataset. **c**, UMAP visualizations of integration results for SpatialFuser, SPACEL, STAligner, and SLAT on slices 1 and 2 of the BaristaSeq dataset, coloured by region labels (top) and batch labels (bottom). **d**, ARI scores before and after integration for domain identification in slices 1 and 2 of the BaristaSeq dataset. **e**, Alignment of Stage 54 and Stage 57 slices from the Stereo-seq axolotl regenerative telencephalon dataset, coloured by region labels (300 alignment pairs shown for clarity). **f**, Sankey plot showing region type correspondence based on alignments from SpatialFuser for Stage 54 and Stage 57 slices of the Stereo-seq dataset. **g**, UMAP visualizations of integration results for SpatialFuser on Stage 54 and Stage 57 slices of the Stereo-seq dataset, coloured by region labels (top) and batch labels (bottom). **h**, Spatial domains in the Stage 57 slice of the Stereo-seq dataset identified by Mclust clustering on SpatialFuser-integrated embeddings. **i**, Expression distributions of FGF17, STRA6, and SLIT2, specifically enriched in a potential subregional structure within the dEGCs region of the Stage 57 slice from the Stereo-seq dataset.

An effective integration method should not only achieve accurate alignment but also preserve underlying biological patterns in the latent space. To this end, we further evaluated the spatial fidelity of the integrated embeddings. UMAP visualization showed that all four methods effectively removed batch effects (Fig. 3c). With original annotations used as supervision, SPACEL maintained strong region-level separation (Supplementary Fig. 4), but its embedding space appeared fragmented (Supplementary Fig. 5), suggesting limited intra-domain coherence despite minimal cross-domain mixing. STAligner and SLAT preserved the hierarchical structure of the six cortical layers to some extent, but layer boundaries were blurred and showed considerable mixing between adjacent layers. Compared to these methods, SpatialFuser produced embeddings with clearer expression patterns showed in UMAP (Fig. 3c & Supplementary Fig. 5) and more hierarchically organized, anatomically consistent domain structures that closely matched the ground truth (Supplementary Fig. 4).

To quantitatively compare the effectiveness of these methods in supporting integrative spatial heterogeneity analysis, we performed clustering on the joint embeddings from each pair of BaristaSeq slices using the Mclust algorithm, focusing primarily on the ARI before and after alignment and integration. Since SLAT and SPACEL operates only in multi-slice mode, ARI scores before integration were left unpopulated. The bar plot (Fig. 3d) shows that SpatialFuser demonstrated the best performance for post-integration, and the integration process enabled more accurate capture of spatial patterns, as clustering on the integrated embeddings yielded spatial domains more consistent with the ground truth (average ARI=0.86).

We further evaluated SpatialFuser’s efficacy in integrating non-consecutive slices from different developmental stages, which exhibit significant differences in cell-type composition and distribution. We employed a Stereo-seq[12] dataset representing axolotl regenerative telencephalon[10], which has manual annotations for each tissue region as reference, and applied SpatialFuser to align and integrate two slices from stage 54 and stage 57 (Fig. 3e). Through alignment, most spots were accurately matched despite substantial variation in spot numbers and types across slices (Fig. 3f), and dynamic changes in cell identities during development were successfully captured (e.g., mature nptxEX at stage 57 could be traced back to immature nptxEX at stage 54). In parallel, SpatialFuser effectively corrected batch effects while preserving expression pattern differences among distinct cell-types (Fig. 3g).

Interestingly, a large region of development-related ependymoglial cells (dEGCs) was enriched with less-aligned cells during the iterative training process of SpatialFuser (Supplementary Fig. 9). We hypothesize that this may be attributed to intra-regional heterogeneity within the seemingly homogeneous dEGCs domain. For instance, previously unannotated substructures may have emerged during telencephalon development from stage 54 to stage 57, reducing cross-slice similarity within this region and thereby hindering the formation of high-quality correspondences. To support this hypothesis, we performed Mclust clustering on the integrated embeddings and identified a distinct subpopulation within the dEGCs domain (Cluster 17) (Fig. 3h), likely reflecting emerging subregional or functional differentiation. Further spatial differential analysis confirmed that this subregion exhibits a unique spatial expression pattern, with FGF17, STRA6, and SLIT2 significantly enriched (Fig. 3i). These genes are known to play key roles in embryonic development and neurodevelopmental processes[41-44], aligning well with the spatial distribution and potential functions of the subregion. Collectively, these findings demonstrate SpatialFuser’s capability to reveal dynamic cellular changes through spatiotemporal integrative analysis.

### Effective and efficient spatial multi-omics integration enables complementary tissue profiling

Incorporating multiple modalities into the analysis of biological samples holds great potential for deepening our understanding of the mechanisms underlying cellular and tissue organization[45]. To evaluate the multi-omics integration capability of SpatialFuser, we applied it to spatial ATAC–RNA-seq data from an embryonic day 13 (E13) mouse embryo section (Fig. 4a), in which the telencephalon region was annotated based on marker genes for the cortical plate, ventricular zone, and striatal primordium (Fig. 4b), and compared its performance with current state-of-the-art methods. Leiden clustering of the integrated embeddings from all models recovered the Eye region (cluster 9 in both RNA and ATAC Leiden clustering) (Fig. 4c). However, only SpatialFuser and SpatialGlue preserved the layered organization of the hindbrain with high structural fidelity. Focusing on the dorsolateral telencephalon in the forebrain, SpatialFuser and MISO uniquely distinguished the developing pallial cortical plate from the pallial ventricular zone. This neuronal population was previously shown to be difficult to resolve using either modality alone, as demonstrated by joint epigenomic and transcriptomic profiling in a prior study [14]. During embryonic telencephalon development, key transcription factors exhibit spatial gradients rather than sharply defined boundaries [17, 46]. To quantify the spatial concordance between identified CP clusters and CP marker enrichment, we evaluated cluster–marker agreement using purity and coverage metrics across a range of marker-enriched spot definitions (see Methods). Although the cortical plate cluster identified by MISO shows near maximal marker purity (Fig. 4d, left), it is largely restricted to localized expression hotspots. In contrast, SpatialFuser delineates a broader and spatially continuous region, maintaining a high level of marker purity while achieving substantially higher coverage score (Fig. 4d, right), consistent with the gradual transitions expected across developing cortical territories. By comparison, MultiGATE produces less distinct regional boundaries despite achieving overall modality alignment (Fig. 4c and Supplementary Fig. 10), whereas SIMO showed limited recovery of the original anatomical domains even when label information is incorporated during training (Fig. 4c), which might be because its original objective differs from direct slice-level spatial interpretation and spot-level integration. We also observed that increasing the number of mapped cells in SIMO leads to increasing spatial disorganization and reduced domain consistency (Supplementary Fig. 11).

**Fig. 4.**
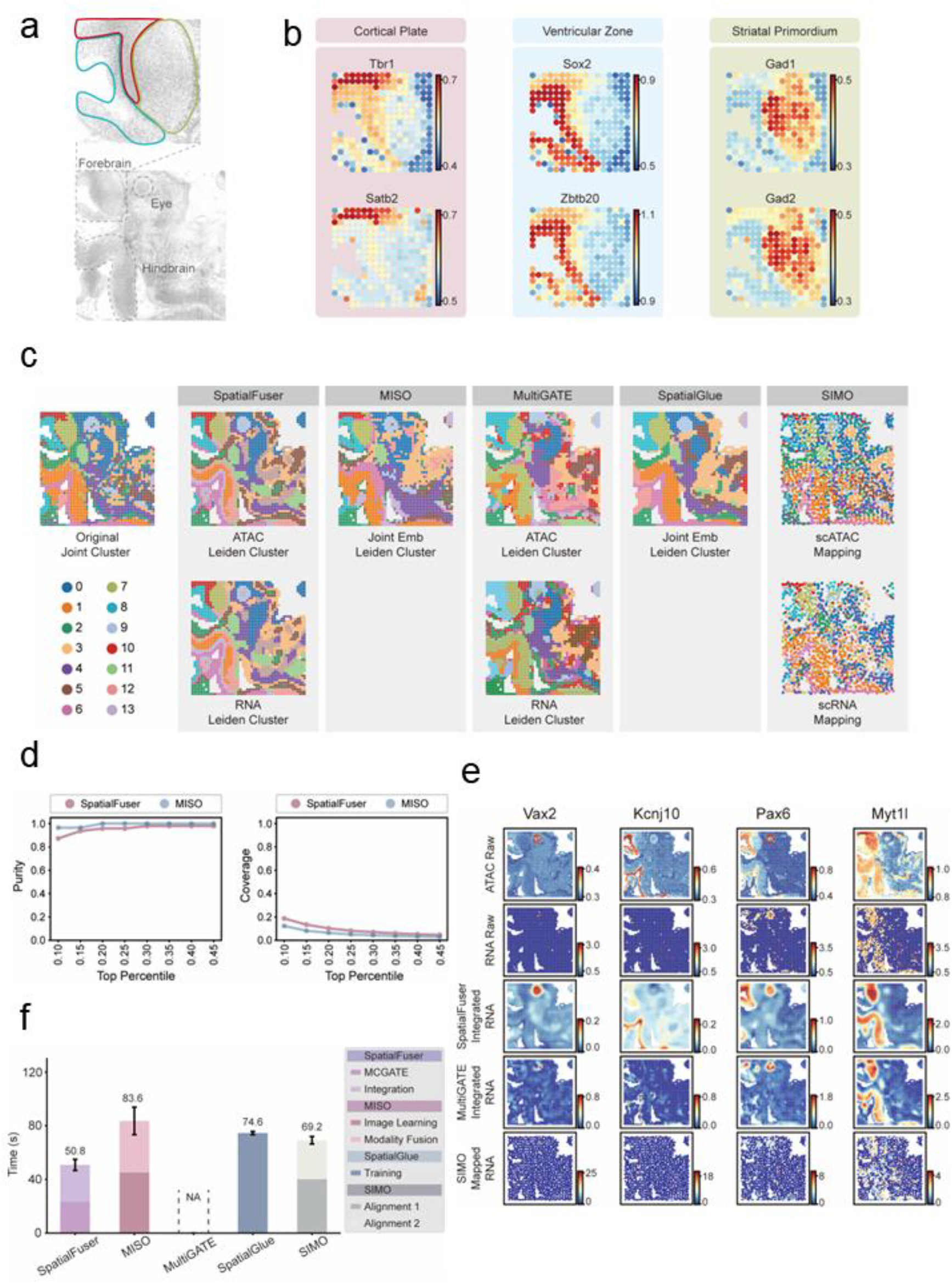
Fast and precise integrative analysis of epigenome–transcriptome co-profiled mouse embryo slice. **a**, H&E-stained histological image of spatial ATAC– RNA-seq data from an E13 mouse embryo, with magnified and annotated views of the telencephalon. **b**, Spatial gene expression patterns of marker genes for the cortical plate, ventricular zone, and striatal primordium in the telencephalon. **c**, Spatial domains identified by Leiden clustering on integrated embeddings or mapped features generated by SpatialFuser, MISO, MultiGATE, SpatialGlue, and SIMO using the E13 mouse embryo spatial ATAC–RNA-seq dataset. SpatialFuser, MultiGATE, and MISO produce modality-specific representations, whereas SpatialGlue and SIMO learn joint representations across modalities. **d**, Purity–Coverage analysis of cortical plate marker enrichment under varying percentile thresholds. **e**, Spatial expression patterns of Vax2, Kcnj10, Pax6, and Myt1l in the original ATAC-derived gene activity, original RNA expression, and reconstructed or mapped RNA profiles produced by SpatialFuser, MultiGATE, and SIMO after integration. **f**, Computational efficiency comparison of methods evaluated on the E13 mouse embryo spatial ATAC–RNA-seq dataset.

At embryonic day 13, chromatin accessibility may not fully resolve all the cell types defined by transcriptional profiles, potentially due to lineage priming[47, 48], as some genes exhibit differing spatial signal patterns between ATAC and RNA modalities. For instance, Vax2 (ventral retina marker[49]), Kcnj10 (ventricular zone and astrocyte progenitor marker[50]), Pax6 (cortical progenitor and optic vesicle marker[51]), and Myt1l (hindbrain neuron marker[52]) displayed relatively low RNA expression in their respective regions at stage E13 in the mouse embryo, despite marked chromatin accessibility. For methods that reconstruct omics features, we further evaluated their ability to capture complementary expression patterns by integrating chromatin accessibility with transcriptomic signals. Compared with MultiGATE and SIMO, SpatialFuser preferentially enhances the detection of weakly expressed yet epigenetically informative genes (Fig. 4e). However, it is important to note that while SpatialFuser functions as an integrative enhancement tool that leverages complementary information from different modalities to reconstruct a more comprehensive and biologically meaningful gene expression landscape, it does not explicitly differentiate whether low RNA expression arises from true lineage priming or technical limitations in transcript detection. Disentangling these causes requires further experimental validation.

As spatial multi-omics datasets continue to increase in scale and complexity, computational efficiency becomes increasingly important for integration methods. We therefore compared training runtime across methods using the E13 mouse embryo spatial ATAC RNA-seq dataset (see Methods), and the results show that SpatialFuser required less training time than all other approaches (Fig. 4f). Of note, MultiGATE is implemented in legacy software environments and built on an early version of the TensorFlow framework, which posed practical challenges for deployment on our GPU systems. Therefore, MultiGATE was trained on CPU, with each run requiring approximately 1.5 to 2 hours. Together, our findings indicate that SpatialFuser can effectively capture molecular signals associated with transcriptional readiness within complex tissue contexts that remained difficult to resolve using unimodal or earlier integration approaches. This capability facilitates the identification of emerging cell subtypes or transitional cellular states relevant to developmental processes and early disease progression.

### Cross-resolution integration of weakly correlated spatial modalities advances tissue-level information

Compared to the one-to-one correspondence between the genome and transcriptome, the linkage between the proteome or metabolome and the transcriptome or epigenome is generally less direct[53], as protein abundance and metabolite levels are influenced by multiple regulatory processes beyond gene expression, such as translation, post-translational modifications, and cellular environment[54, 55]. To explore cross-modality integration under such weakly correlated conditions, we applied SpatialFuser, with NDT-based coordinate pre-matching, to integrate and align transcriptomic (MAGIC-seq[11]), proteomic (PLATO[15]), and metabolomic (MALDI-MSI[7]) data under varying resolutions and mismatched tissue geometries obtained from consecutive mouse cerebellum tissue slices. Result shows SpatialFuser effectively captured shared spatial molecular patterns across different omics layers and successfully corrected modality biases while preserving true biological variation, as evidenced by the spatial domains identified through Louvain clustering of the latent embeddings, which aligned well with the original anatomical annotations (Fig. 5a). Most spots across the tissue were well aligned, with high alignment similarity scores (Supplementary Fig. 13) and matching accuracy (Supplementary Fig. 14, Transcriptome–Proteome: 0.84, Transcriptome–Metabolome: 0.75, Proteome–Metabolome: 0.86). It is noteworthy that manual annotation is affected by the granularity and distribution discrepancies across modalities. For example, although MAGIC-seq transcriptomic data and PLATO proteomic data share the same spatial coordinates, the matching rate of their original labels is only 83%, which undermines the reliability of using label-matching accuracy as a metric to evaluate spot-to-region mapping performance. By visualizing the embeddings with UMAP, we further demonstrate SpatialFuser’s ability to accurately align the same cell types across different omics modalities, as spots of the same type are consistently co-localized in the embedding space, even under weak cross-modality cellular correspondence (Fig. 5b).

**Fig. 5.**
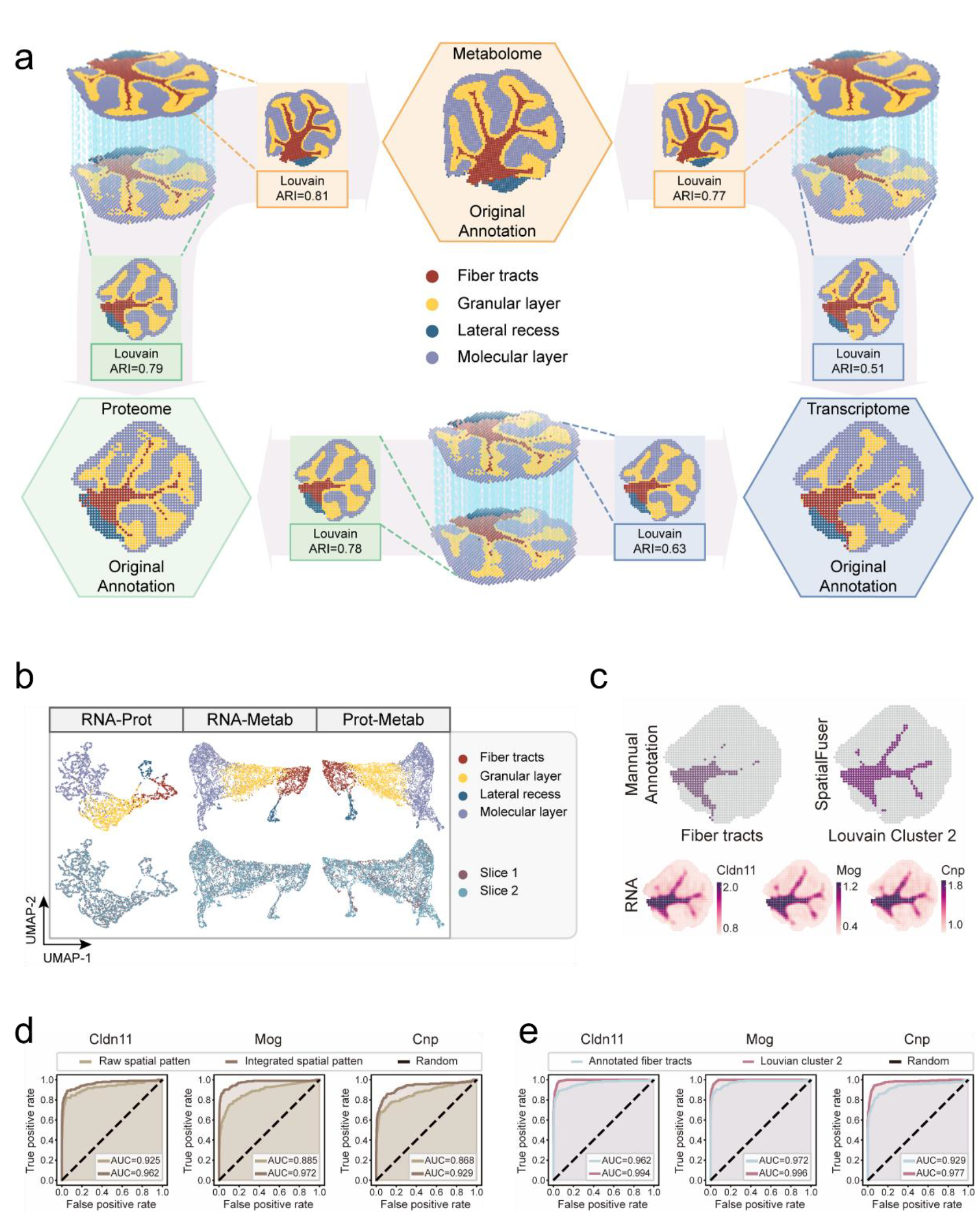
SpatialFuser effectively integrates and aligns cross-resolution datasets from consecutive mouse cerebellum tissue slices across transcriptomic, proteomic, and metabolomic data. a, Pair-wise integration and alignment (300 alignment pairs shown for clarity) of transcriptomic (MAGIC-seq), proteomic (PLATO), and metabolomic (MALDI-MSI) data from consecutive mouse cerebellum tissue slices, with Louvain clustering for tissue domain detection. b, UMAP visualizations of modality bias correction results from SpatialFuser on pairwise integrations of the mouse cerebellum datasets, coloured by region labels (top) and batch labels (bottom). c, Distribution of the original Fiber tract annotation in the MAGIC-seq slice and the corresponding RNA–metabolome integration result (Louvain cluster 2), alongside spatial expression patterns of the Fiber tract marker genes Cldn11, Mog, and Cnp. d, ROC curves of marker gene expression before and after integration for distinguishing the original Fiber tract annotation. e, ROC curves of reconstructed marker gene expression distinguishing Louvain-detected Fiber tracts (cluster 2) and the original Fiber tract annotation.

Due to differences in spatial resolution and barcode density, the MALDI-MSI slice captured 3908 spots, which is significantly more than the 1677 spots in the transcriptomic and proteomic datasets, thereby enabling finer-grained tissue domain annotations. Through cross-modality and cross-resolution integration and alignment, SpatialFuser is able to leverage the detailed histological distribution patterns in the metabolomic data to enhance the delineation of fiber tracts in the spatial transcriptomic data, which were previously difficult to resolve (Fig. 5c). This refined delineation may reduce agreement with the original coarse annotation-based metrics, but it provides a biologically testable hypothesis that can be evaluated using marker gene concordance. To assess whether the reconstructed transcriptomic profiles preserved biologically relevant spatial patterns, we evaluated the ability of known canonical oligodendrocyte marker genes (Cldn11[56], Mog[57], and Cnp[58]) to discriminate the originally annotated fiber tracts before and after integration. ROC curve analysis showed that the reconstructed features achieved higher AUC scores in distinguishing the fiber tracts defined by the original annotation (Fig. 5d), indicating improved concordance with expected spatial expression patterns. Notably, when using Louvain-identified Fiber tracts cluster as the reference, we observed even higher AUC scores (Fig. 5e). The strong spatial concordance between the reconstructed marker gene expression patterns and the refined anatomical delineation supports the biological validity of the finer-grained structure revealed by the SpatialFuser integration.

Together, these results demonstrate the versatility of SpatialFuser in decoding complex biological systems through spatial multi-omics integration. By bridging modalities with different biological meanings and technical characteristics, SpatialFuser enables deeper and more comprehensive tissue profiling across resolutions, offering a powerful tool for spatial systems biology.

## Discussion

Spatial technologies are expanding into multi-omics, enabling researchers to measure transcriptomics, epigenomics, proteomics, metabolomics, and other molecular layers at defined tissue locations. Underlying this is an unprecedented opportunity to integrate and interpret molecular features from diverse complementary perspectives for a deeper understanding of biological mechanisms within tissue microenvironment[59-61]. In this study, we propose SpatialFuser, a deep learning-based framework designed for detailed molecular profiling within individual tissue sections and cross-modality, multi-sample integrative analysis of spatial multi-omics data. Through benchmarking on real spatial multi-omics datasets generated by diverse technologies, we show that SpatialFuser outperforms existing state-of-the-art methods in spatial domain detection, consecutive slice alignment, and spatial multi-omics data integration. Notably, we further demonstrate SpatialFuser’s unique capability for cross-resolution integration of weakly correlated modalities, which has not been previously established. By combining flexible single-slice representation learning with coupled matching–fusion optimisation for cross-slice analysis, SpatialFuser provides a coherent solution for accurate spatial interpretation, robust cross-slice alignment, and effective cross-modality integration of unpaired spatial multi-omics data. To accommodate diverse input modalities, SpatialFuser introduces MCGATE to learn unified representations from multi-platform spatial omics data. Unlike methods such as STAGATE and SEDR that primarily focus on local information propagation, MCGATE employs a multi-head collaborative attention mechanism that further captures long-range feature correspondences in the embedding space, enabling more accurate identification of spatially coherent patterns. The effectiveness of this design is supported by ablation and interpretability analyses, and beyond performance gains, the architecture improves model transparency while providing new perspectives for interpreting spatial molecular organization.

SpatialFuser introduces methodological advances and broader utility in cross-slice alignment and multi-modality integration compared with previous approaches. Unlike alignment methods such as STAligner and SPACEL, SpatialFuser learns slice-specific embeddings that preserve native feature structures and avoid information loss from forced dimensionality matching, while applying a more relaxed alignment strategy to model cross-slice correspondences, thereby better supporting scenarios involving different technologies. Beyond alignment, integration tools such as MISO, SpatialGlue, and MultiGATE are designed for co-profiled data measured on the same tissue section and often assume similar cellular composition or strong biological priors. Rather than relying on one-to-one molecular correspondences across modalities, SpatialFuser captures correlated spatial molecular patterns in the embedding space to correct batch effects and modality biases. It learns latent correspondences directly from data and is built to handle unpaired sections across different samples, resolutions, technologies, and developmental stages, extending multi-omics analysis to a broader range of real-world scenarios.

A current limitation of SpatialFuser lies in the memory footprint of multi-head collaborative attention mechanism. Although multi-head attention improves representation learning by aggregating feature relationships across multiple subspaces, the lack of memory-efficient parallel operations for high-dimensional sparse matrices in current deep learning frameworks leads to substantial GPU memory demands for large datasets [62, 63]. To improve scalability, SpatialFuser also provides a sparse-optimized single-head mode, which can be used for large-scale tissue sections when computational efficiency is prioritized. We anticipate that ongoing advances in hardware and software will mitigate these bottlenecks. Moreover, we plan to develop more efficient sparse computation strategies in future versions of SpatialFuser to better support analyses of increasingly large tissue sections at higher spatial resolution. In fact, SpatialFuser remains generally fast. MCGATE embedding generation requires only ~30 seconds per DLPFC slice on an NVIDIA GeForce RTX 4090 GPU (24 GB), and integration efficiency is reflected in our benchmarking results.

Although image-derived features can improve tissue region identification and spatial molecular pattern recognition[25, 64], SpatialFuser does not currently include a dedicated histology embedding module, as histological images are not consistently available across spatial omics platforms, limiting the general applicability of image-dependent models. Moreover, effective image representation learning often relies on large neural networks with substantial training costs. In contrast, SpatialFuser is intentionally lightweight. In lower-throughput modality analysis scenarios such as CODEX, SpatialFuser may involve only a few thousand trainable parameters, which supports its deployment in resource-constrained settings.

In summary, SpatialFuser does not merely incrementally improve upon existing methods, but also provides a versatile, robust, and broadly applicable framework that empowers researchers to integrate spatial multi-omics data across a wide spectrum of experimental designs, beyond the limitations of current methods. To facilitate broad adoption by the research community, we have released the SpatialFuser package along with detailed tutorials and demo cases for all presented experiments at our GitHub repository https://github.com/liwz-lab/SpatialFuser.

## Methods

### Data Preprocessing

#### Filtering and normalization

SpatialFuser takes molecular feature matrices and spatial coordinates from spatial multi-omics data as input. For consistency across platforms, we refer to both cells and capture voxels in spatial omics experiments as spots. Spots lacking valid annotations were excluded from analyses that required ground-truth labels or quantitative evaluation. To make coordinate scales comparable across slices, the spatial coordinates of each spot were normalized to [0,1]. For each slice, we normalized the raw molecular feature matrix by total counts across all features, scaled each spot to 10,000 total counts, and applied log-transformation to stabilize variance and reduce the influence of extreme values. For high-throughput data, including sequencing-based transcriptomics, ATAC-derived gene activity profiles, and AI-enhanced proteomics, we selected the top 3,000 spatially variable features. For lower-throughput modalities, including image-based transcriptomics, CODEX proteomics, and MALDI-MSI metabolomics, all features were retained. Normalization and feature selection were conducted using Scanpy[65] package.

#### Spatial graph construction

We denote a spatial omics slice as *D* = {(*x*_*i*_, *s*_*i*_), *i* = 1,2, ⋯, *N*}, where *N* is the number of spots within the slice, *x*_*i*_ ∈ ℝ^*d*^ is the preprocessed feature vector of spot *i, s*_*i*_ ∈ ℝ^2^ is its spatial coordinate, and *d* is the number of omics features. SpatialFuser constructs a spatial graph by connecting each spot to its K-nearest neighbours (KNN) in the coordinate space. The neighbour set of spot *i* is denoted as *N*_*i*_. The value of K is selected according to spatial domain heterogeneity and connectivity patterns across different multi-omics scenarios and platforms.

### MCGATE for multi-omics data

#### Encoder

Given input *D*, for spot *i*, the *k*-th encoder layer (*k* ∈ {1,2, ⋯, *L*}) adaptively learns the pairwise relationship between spot *i* and each neighbouring spot *j* ∈ *N*_*i*_ through an edge attention mechanism. For each neighbouring spot *j*, the edge attention score 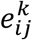 at layer *k* is first computed as:

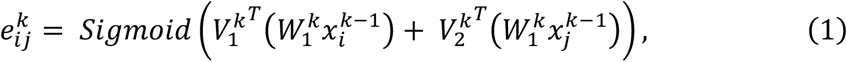

where 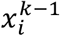 and 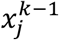 denote the representations of spot *i* and *j* from the previous layer, *W*_1_ is a learnable weight matrix, and *V*_1_ and *V*_2_ are trainable weight vectors. For spot *i*, the edge attention scores over its neighbour set *N*_*i*_ are then normalized using a sparse *Softmax* function to obtain the attention weights for information aggregation:

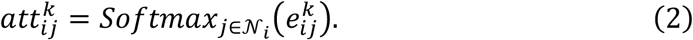

Based on these attention weights, the encoder aggregates information from neighbouring spots in *N*_*i*_ to compute the neighbourhood-aware representation 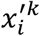:

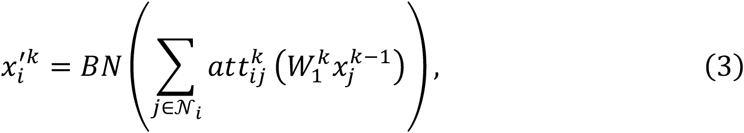

where *BN* denotes batch normalization[66], which stabilizes training and accelerates convergence[67].

Finally, a feedforward transformation with residual connection is applied to introduce nonlinearity and enhance the model’s representational capacity:

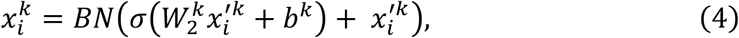

where *W*_2_ is a learnable weight matrix, *b* is a trainable bias, and *σ* denotes the activation function. The output of the final encoder layer 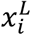 serves as the low-dimensional feature embedding learned by MCGATE.

#### Adaptive adjacency expansion layer

Adaptive adjacency expansion layer. Previous spatial graph methods commonly prioritize local neighbourhoods for information aggregation, reflecting the local coherence and spatial contiguity of tissue organization. However, spatial omics data may also contain molecularly similar spots across non-adjacent tissue regions, which cannot be fully captured by purely local aggregation. Directly computing global attention over all spot pairs in the high-dimensional input space is computationally expensive and scales poorly with the number of spots. To address this, we introduce a collaborative attention mechanism that preserves neighbourhood information while selectively incorporating long-range signals from molecularly similar but spatially distant spots in a low-dimensional embedding space, allowing MCGATE to model multi-scale spatial expression patterns efficiently.

The adaptive adjacency expansion layer first computes pairwise cosine similarity between final-layer spot embeddings to derive local and long-range feature-similarity patterns:

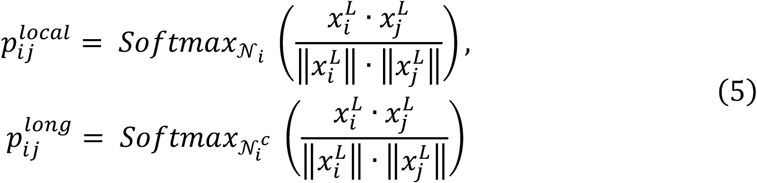

where *N*_*i*_ denotes the spatial neighbourhood of spot *i*, and *N*_*i*_^*c*^ denotes spots outside this spatial neighbourhood.

The local similarity pattern 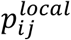 is fed back to the encoder to enhance the local attention mechanism previously defined in Equation (2):

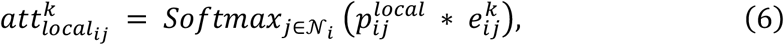

where 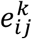 is the edge attention score computed at *k*-th encoder layer.

For each spot *i*, the adaptive adjacency expansion layer selects mutual nearest neighbour (MNN) spot pairs[68] with the highest similarity from 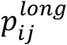, generating a sparse long-range attention graph for controlled non-local information aggregation. During training, let *G*_*i*_ denotes the set of long-range neighbours for spot *i*, the long-range attention weights are computed as:

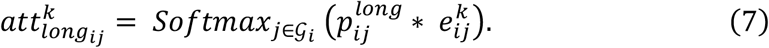

The collaborative attention weights are then defined a weighted sum of local and long-range attention:

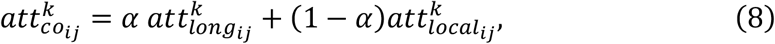

where *α* ∈ [0,1] (default *α* = 0) is a slice-specific hyperparameter that controls the contribution of long-range attention during information aggregation. By integrating local information propagation with controlled non-local aggregation, the adaptive adjacency expansion layer enhances information flow across spatially adjacent and molecularly similar but spatially distant spots, thereby improving MCGATE’s ability to capture multi-scale spatial expression patterns.

#### Decoder

Analogous to the encoder, the *k*-th decoder layer (*k* ∈ {1,2, ⋯, *L*}) reconstructs the representation of spot *i* as follows:

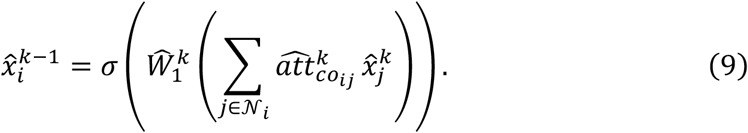

The decoder takes the low-dimensional feature embedding x^L^ as input and outputs the reconstructed feature representation 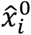, which approximates the pre-processed input feature vector 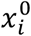.

To reduce the risk of overfitting, MCGATE adopts a parameter sharing strategy between the encoder and the decoder. Specifically, for the *k*-th decoder layer, we set 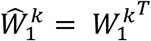 and 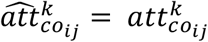, thereby sharing the transformation matrices and attention weights. However, the decoder does not replicate the feedforward block of the encoder, thereby avoiding overly rigid architectural symmetry.

The training of MCGATE is driven by a reconstruction objective that minimizes the mean squared error (MSE) between the pre-processed input features and decoder output:

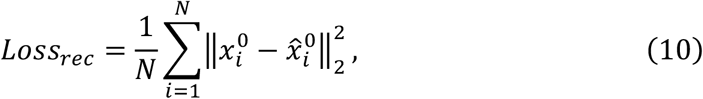

where *N* is the total number of spots. As 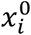 denotes the normalized and log-transformed input feature vector, MSE is applied to transformed continuous features rather than raw counts. This reduces the dominance of extreme values and provides a simple modality-agnostic reconstruction objective across heterogeneous omics data. However, MSE does not explicitly model zero inflation or distinguish biological absence from technical dropout, and such distinctions require modality-specific probabilistic modelling because sparsity levels can differ substantially across technologies and modalities.

#### Multi-head attention

Inspired by previous work[69], MCGATE incorporates multi-head attention to capture feature relationships across multiple representation subspaces. Specifically, the input features are projected into *H* representation subspaces, each corresponding to one attention head. In the *k*-th encoder layer (*k* ∈ {1,2, ⋯, *L*}), the *h*-th collaborative attention weight (*h* ∈ {1,2, ⋯, *H*}) between spots *i* and *j*, denoted 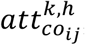, is computed in parallel using head-specific parameters 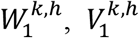, and 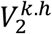. In the adaptive adjacency expansion layer, cosine similarity is also computed separately for each head, allowing each subspace to define its own local and long-range feature-similarity patterns.

The outputs of all attention heads are concatenated to form the multi-head representation. The encoder aggregation formula, extending Equation (3), is defined as:

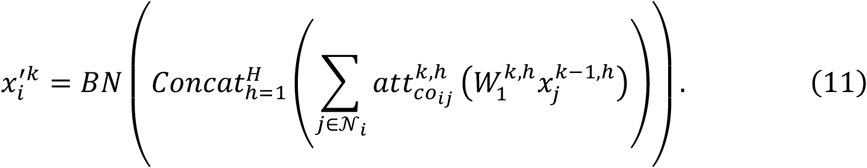

Similarly, the decoder reconstruction formula, extending Equation (9), becomes:

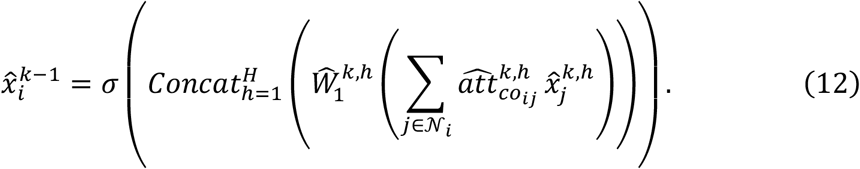

Due to limited support for efficient multi-head sparse tensor operations in current deep learning frameworks (e.g., PyTorch[69]), the multi-head attention mechanism can impose substantial GPU memory demands. To improve scalability for large-scale spatial multi-omics datasets, MCGATE provides a sparse-optimized single-head mode that reduces memory usage and computational cost during training.

## Matching layer

SpatialFuser models the alignment of spatial omics slices as a graph matching problem and solves it through optimal transport. The matching layer aims to estimate an optimal coupling matrix that defines cross-slice correspondences by minimizing the Wasserstein distance[70] between embedding distributions. It operates on slice-specific MCGATE embeddings projected by the shared fusion layer and provides candidate anchor pairs for subsequent fusion. Although these initial fused embeddings are represented in a common latent space, they are not assumed to be fully aligned. We therefore introduce geometric constraints to guide early matching, with embedding similarity serving as auxiliary representation-level evidence rather than a direct distance comparison between independent embeddings.

Let *E*_1_ ∈ ℝ^*n*×*d*′^ and *E*_2_ ∈ ℝ^*m*×*d*′^ denote the fused embeddings of two slices, where *n* and *m* are the numbers of spots and *d*′ is the embedding dimension. SpatialFuser defines a geometry-constrained cross-slice similarity measure as follows:

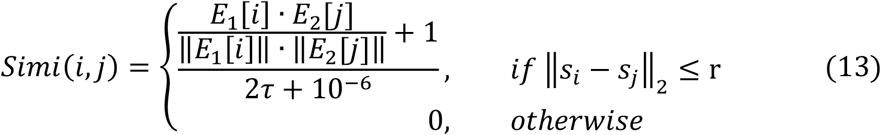

where the indices *i* ∈ {1,2, ⋯, *n*} and *j* ∈ {1,2, ⋯, *m*} denote spots from the two slices, s_i_ and *s*_*j*_ represent their corresponding normalized spatial coordinate vectors, *τ* is a temperature coefficient, and r denotes the spatial neighbourhood radius. The radius r defines a geometry-constrained candidate window for early cross-slice matching in the normalized coordinate space.

Based on the similarity matrix defined in Equation (13), SpatialFuser formulates the alignment task as the following optimal transport problem:

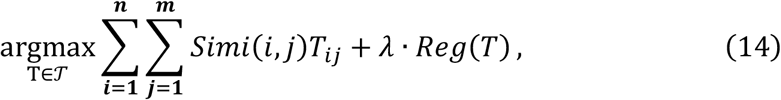

where *T* ∈ [0,1]^*n*×*m*^ is the soft coupling matrix to be optimized. The set *T* = {*T* ∈ ℝ^*n*×*m*^ |*T*1_*m*_ ≤ 1_*n*_, *T*^*T*^1_*n*_ ≤ 1_*m*_} defines the feasible region. *Reg*(⋅) denotes the entropy regularizer, and λ is its weighting factor. SpatialFuser employs the Sinkhorn algorithm[71] to solve this entropic optimal transport problem through efficient iterative updates.

Although Equation (14) estimates cross-slice correspondences over the original spot-to-spot similarity matrix, heterogeneous spatial omics datasets may contain spots without reliable counterparts because slices can differ in cell-type composition, spatial resolution, and spot number. To accommodate unmatched spots and ambiguous assignments, SpatialFuser introduces a relaxed matching strategy by augmenting the original similarity matrix *Simi* ∈ ℝ^*n*×*m*^ defined in Equation (13) with additional dustbin channels, allowing low-confidence assignments to be absorbed by the dustbin rather than forced into spot-to-spot matches:

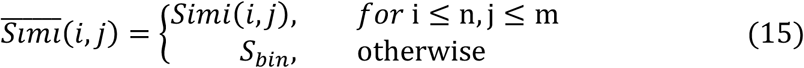

where *i* ∈ ℤ_[1,*n*+1]_ and *j* ∈ ℤ_[1,*m*+1]_, 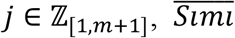 is the augmented similarity matrix with the dustbin channels, and *S*_*bin*_ is a constant weight (*S*_*bin*_ = 1 in default) assigned to the dustbin.

The dustbin channel strategy relaxes the original matching problem by allowing the augmented similarity matrix to be optimized under an equality-constrained transport formulation:

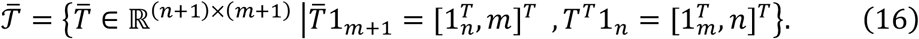

By allowing ambiguous or low-confidence assignments to be absorbed by the dustbin channels, SpatialFuser reduces forced spot-to-spot alignment under compositional or resolution mismatch.

To provide sufficient anchor pairs for fusion layer during early training and further improve matching quality, SpatialFuser introduces a Mutual Top-K Neighbour (MKN) strategy based on the soft assignment matrix generated by the Sinkhorn algorithm. Specifically, the model first selects MNN pairs from the soft couplings as high-confidence correspondences. However, batch effects or modality bias in the early training may result in insufficient MNN pairs. To address this, the matching criterion is relaxed to include mutual top-K neighbours, providing additional anchor pairs for the fusion layer and stabilizing training. During iterative optimisation, these filtered anchors are updated after fusion-layer refinement, allowing initial coarse correspondences to be refined as the latent space becomes progressively more comparable.

### Fusion layer

In the fusion layer, a shared multilayer perceptron (MLP) is connected to two independent MCGATE modules to project the low-dimensional embeddings of spatial omics data from different platforms or modalities into a unified latent space. To reduce the risk of learning batch-specific ordering patterns, the embeddings are concatenated and randomly permuted along the spot dimension before being passed into the MLP.

The fusion layer employs contrastive learning for spatial multi-omics data integration. Specifically, high-confidence cross-slice correspondences derived from the matching layer are used as anchor-positive pairs to guide the correction of batch effects or modality bias. Negative samples are randomly drawn from non-neighbouring regions within the same tissue slice to improve cell-type discriminability while reducing the probability of sampling false negatives. Because the number of spatial spots is typically much larger than the local neighbourhood size, this strategy provides an efficient approximation for negative sampling.

The training process employs an improved triplet loss[31] as the modality fusion objective, which pulls anchor-positive pairs closer, separates anchor-negative pairs, and enforces both absolute and relative distance constraints to improve training stability:

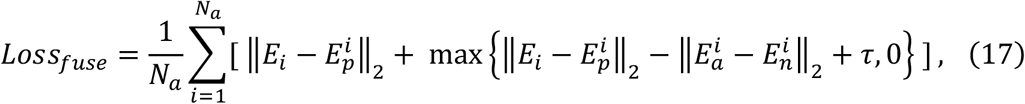

where *E*_*i*_ is the fused embedding of spot *i*, serving as anchor, 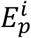 and 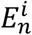 denote the embeddings of the corresponding positive and negative spots respectively, and *τ* is a margin parameter. To prevent embedding collapse and preserve feature representational fidelity, a reconstruction loss is incorporated into the integration process. In addition, to mitigate potential inconsistencies caused by the dual independent encoder design of SpatialFuser, we introduce an embedding direction constraint loss *Loss*_*dir*_ to align the global orientation of the representations across slices:

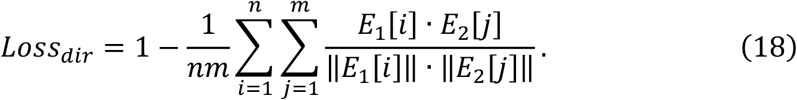

The total objective function of the integration process combines the fusion objective, the reconstruction loss, and the embedding direction constraint loss, and is formulated as:

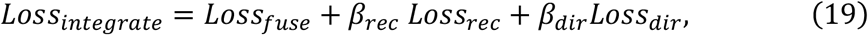

where *β*_*rec*_ and *β*_*dir*_ are the weights of the reconstruction loss and the embedding direction constraint loss (*β*_*rec*_ = 1, *β*_*dir*_ = 0.1 in default), respectively. The embeddings integrated by the fusion layer are used in the next iteration to update the matching layer, enabling cross-slice correspondences and shared representations to be refined jointly across training rounds. This iterative refinement reduces dependence on initial matches derived from incompletely aligned embeddings and mitigates error propagation during matching–fusion.

### Coordinate pre-matching module (optional)

Spatial registration of spot coordinates is an important preparatory step for aligning and integrating spatial omics slices when substantial geometric mismatch exists. SpatialFuser represents spot coordinates as 2D point clouds and reduces coordinate discrepancies using rigid transformations defined by a rotation matrix *M* and a translation *t*. Given a source point cloud *Y*_s_ ∈ ℝ^*n*×2^ and a target point cloud *Y*_t_ ∈ ℝ^*m*×2^, where *n* and *m* are the numbers of spots in each slice, coordinate pre-matching estimates a rigid transformation Φ_M,t_ that maps the source coordinates to the target coordinate space:

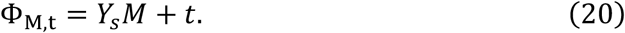

The transformation parameters are optimized by minimizing a registration loss defined by the selected registration algorithm:

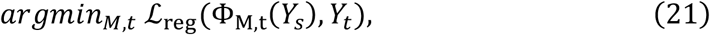

where ℒ_reg_ denotes the algorithm-specific registration optimization objective.

SpatialFuser provides multiple registration options to accommodate different geometric scenarios. The ICP algorithm is a classical method for solving point cloud registration problems[72, 73] and has been applied to spatial omics pre-registration when slices have relatively regular spatial layouts or comparable tissue boundaries. It estimates spot correspondences by nearest-neighbour search between slices and iteratively update the rotation matrix *M* and translation vector *t* by minimizing the Euclidean distance between matched spot pairs. In our implementation, ICP uses alpha-shape boundary points extracted from the spot coordinates, updates correspondences automatically by nearest-neighbour search, and optimizes the rigid transformation without manually selected landmarks.

As an alternative registration option, SpatialFuser also incorporates the NDT algorithm[74], which models the reference slice by partitioning it into a 2D grid. Each grid cell *V*_*i,j*_ contains the spots within its bounds, and a multivariate Gaussian distribution *N*(*μ*_*i,j*_, Σ_*i,j*_) is fitted to the spots in each non-empty grid cell. Instead of relying on pairwise Euclidean distances, NDT evaluates the Mahalanobis distance between each transformed spot *z*_*t*_ ∈ ℝ^2^ and the Gaussian distribution of the corresponding grid cell in the reference slice. The coordinate pre-matching is achieved by minimizing the cumulative deviation:

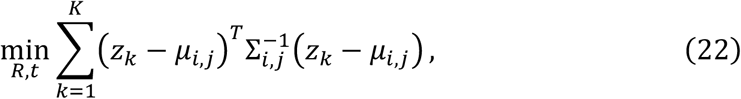

where *K* is the number of the transformed spots used in the NDT objective. This probabilistic approach enables more robust registration in scenarios involving sparse, noisy, or irregular spot distributions[75].

### Whole-slice Alignment

We propose a whole-slice alignment strategy to support downstream analysis and comparison with existing alignment methods. By default, the slice with fewer spatial spots is used as the source slice and is mapped to the slice with more spots. For each source spot *i*, SpatialFuser first identifies a candidate set in the target slice by selecting target spots within the spatial radius *r*, and then computes the cosine similarity between the fused embedding of spot *i* and those of candidate target spots. Inspired by previous work[24], we introduce a probabilistic quality-control procedure to filter unreliable assignments. Instead of randomly pairing spots across both slices, we randomly sample 1,000 spots from the source slice and, for each sampled spot, randomly select a nearby target spot within radius *r* to compute a background similarity score, forming a stricter null distribution. The 95th percentile of this null distribution is used as the quality-control threshold, and the candidate with the highest similarity score above this threshold is retained as the final alignment result for each source spot.

### Implementation details of SpatialFuser

#### Spatial domain detection task

To accommodate datasets with varying complexity, SpatialFuser adopts different MCGATE configurations. For high-throughput data such as 10X Visium, Stereo-seq, and MAGIC-seq, a two-layer MCGATE is used, with hidden dimensions set to 512 and 32 respectively. For low-throughput data such as CODEX and MALDI-MSI, a single-layer MCGATE with 32 hidden units is employed. The 32-dimensional latent space was selected as a compact default representation to balance input feature diversity and computational efficiency across heterogeneous spatial omics datasets. The MCGATE is trained using the Adam optimizer, with the ELU function selected as the activation. The model is trained for 500 steps by default, during which the Adaptive Adjacency Expansion Layer is updated every 100 epochs to refine long-range relationships and their associated attention weights. Both the learning rate and the long-range attention coefficient *α* are customized for each dataset.

#### Alignment and integration task

The training process is divided into two stages: pretraining and contrastive learning. In the pre-training stage, a fusion layer is connected to two independent MCGATE encoders and jointly trained with a reconstruction loss for 500 epochs by default, producing high-quality embeddings for downstream alignment and integration. For slices with semantically similar molecular profiles, shared features across slices are used. The matching layer and the fusion layer are then iteratively optimized to align the embeddings into a shared latent space. Specifically, the matching layer first identifies anchor-positive pairs based on the pre-trained embeddings. These identified pairs are then used to guide the fusion layer in integrating the modalities. By default, triplet pairs are updated every 20 epochs, and this iterative process continues until 500 steps. The learning rate is dataset-specific defined.

#### Hyperparameter selection guideline

Because SpatialFuser handles heterogeneous spatial omics datasets, a single universal default configuration is not technically appropriate. All parameter settings used in this study are provided in the tutorials, and recommended settings according to data characteristics and assay types are summarized in Supplementary Table 1.

#### Clustering, differential expression analysis, and spatial trajectory inference

Clustering was performed by constructing a shared nearest neighbour graph based on the spatially resolved data, followed by cluster identification using the Louvain[35], Leiden[34], or Mclust[33] algorithms. The default resolution parameter used in benchmarking for the Louvain and Leiden clustering algorithms was set to 0.1, while for other downstream analyses it was adjusted to the optimal value for each case. Cluster biomarkers and differentially expressed genes between groups were determined using the Wilcoxon rank-sum test, as implemented in the rank_genes_groups() function of the Scanpy[65] package. To infer spatial trajectories, we applied partition-based graph abstraction via the paga() function in Scanpy.

### Spatial distribution detection across diverse experimental platforms and modalities

#### Data description

The 10X Visium DLPFC dataset[32] comprises 12 tissue sections from 3 individuals, providing whole-transcriptome coverage with 33,438 gene features. Each section contains between 3498 and 4789 spatially resolved spots. The mouse somatosensory cortex dataset[2], generated using the osmFISH platform, includes 5328 spots at single-cell resolution, based on the targeted detection of 33 genes. The CODEX human bladder cancer dataset[4] consists of 75 tumour tissue sections from 31 patients, profiling 35 protein markers across nuclear and membrane compartments. After quality control, ~360,000 epithelial cells, ~140,000 immune cells, and ~90,000 stromal cells were retained. A 1440-dimensional feature matrix was initially constructed using statistical summaries (e.g., percentages and quantile distributions) of protein expression. For the detection of cell type spatial distributions, the 210308_TMA2_reg6 slice was selected, and the average nuclear and membrane expression levels of each protein were extracted as input features. All datasets include detailed annotations of tissue regions or cell types, which serve as ground truth for evaluation.

#### Benchmarking scope

We used the 12-section 10X Visium DLPFC dataset as the primary quantitative benchmark since most methods provide validated workflows for this dataset. The osmFISH and CODEX datasets were used as cross-platform and multi-omics applicability tests rather than exhaustive benchmarks, because most representation learning methods were specifically developed for spatial transcriptomics, whereas multi-omics tools usually require paired modalities and cannot directly support single-slice single-modality analysis.

#### Hyperparameter settings

The hyperparameters for SpatialFuser were optimized through grid search (Supplementary Table 2). For all baseline methods, we adopted the recommended data pre-processing steps and hyperparameter settings as specified in their official implementation and documentation for DLPFC dataset (Supplementary Table 3), except for STAligner. Since STAligner’s embedding model structure is very similar to that of STAGATE, we referred to the documentation of STAGATE for its parameter settings. The documentation links for these methods are shown as follows:

- SpaGCN: https://github.com/jianhuupenn/SpaGCN/blob/master/tutorial/tutorial.md
- STAGATE: https://stagate.readthedocs.io/en/latest/index.html
- SEDR: https://sedr.readthedocs.io/en/latest/index.html

#### Hyperparameter robustness

We tested SpatialFuser’s robustness to the key hyperparameters of MCGATE including (Supplementary Fig. 3): (1) number of encoder and decoder layers, (2) dimension of layers, (3) dimension of MCGATE embedding. We ran MCGATE on all 12 slices of DLPFC dataset and every experiment was run with different random seeds. To identify a suitable attention architecture for the DLPFC dataset, we further benchmarked single-head and multi-head configurations based on the selected network structure.

### Spatiotemporal alignment and integration

#### Data description

The BaristaSeq mouse visual cortex dataset[9] comprises three tissue slices with similar domain distributions and morphological structures, containing 4491, 3545, and 3390 spatial spots, respectively. A total of 78 transcript features were retained, along with detailed anatomical region annotations. Slices 1 and 2 were selected for integration and alignment, with 1,525 and 2,042 annotated spots retained after filtering, respectively. The Stereo-seq axolotl regenerative telencephalon dataset[10] includes 18 brain tissue slices with single-cell resolution, capturing over 20,000 genes across various stages of regeneration and development. Two slices from developmental stages 54 and 57, containing 2929 and 4410 spatial spots respectively, were selected for alignment and integration. The original annotations were used as references for evaluation.

#### Hyperparameter settings

The hyperparameters for SpatialFuser were optimized through grid search, while the hyperparameter settings for all other methods followed the official guidelines and tutorials provided by their respective authors:

- SLAT: https://slat.readthedocs.io/en/latest/tutorials.html
- SPACEL: https://spacel.readthedocs.io/en/latest/index.html
- STAligner: https://staligner.readthedocs.io/en/latest/index.html

To ensure a fair comparison of the integration and alignment performance on consecutive BaristaSeq slices, we conducted a grid search for several key parameters related to both graph construction and network architecture, which are critical yet not explicitly specified in the original documentation of the compared methods. Specifically, SPACEL constructs the spatial graph using 8-nearest neighbours and employs a two-layer spline, with a hidden layer size of 256 and an output embedding dimension of 16 (Supplementary Fig. 6). STAligner builds the graph using 20-nearest neighbours and adopts a two-layer encoder with node dimensions set to 64 and 30, respectively (Supplementary Fig. 7). For contrastive learning-based integration process, the number of nearest neighbours used when constructing MNNs was set to 10. Notably, while typical GCNs tend to suffer from over-fitting, gradient vanishing, or over-smoothing beyond 4–5 layers, SLAT consistently demonstrated improved performance with increased depth, achieving its best results with over 20 LGCN layers (Supplementary Fig. 8). Therefore, we report SLAT’s performance using 20 layers GCN with mlp_hidden = 128 and hidden_size = 64 in the experiments on the BaristaSeq dataset. Before training, SLAT constructs a KNN graph with K=10.

### Spatial ATAC-RNA-seq data integration

#### Data description

The spatial ATAC-RNA-seq dataset[14] of the E13 mouse embryo comprises 2187 spots with genome-wide co-profiling of epigenome and transcriptome. The transcriptomic assay measures 15,748 genes, while the epigenomic assay includes 32,437 accessible chromatin peaks. Gene-level chromatin accessibility was further summarized into activity scores for 24,017 genes derived from the ATAC signal. Manual annotation of anatomical regions was not provided by the authors. Instead, the dataset includes eight major ATAC-defined clusters, 14 RNA-defined clusters, and 14 integrative clusters based on joint analysis of spatial ATAC and RNA data.

#### Hyperparameter settings

For the high-throughput spatial ATAC-RNA-seq data, we constructed a 4-nearest neighbour graph using the top 3000 spatially variable features, which were normalized and log-transformed before being used as input. The number of layers in MCGATE was set to two, with hidden dimensions of 512 and 32, respectively. We first pre-trained two 4-head MCGATE model separately on the ATAC and RNA modalities for 500 epochs using a learning rate of 1e-3 and 1e-4. Further, we set the reconstruction loss weight *β*_*rec*_ = 50 and the directional loss weight *β*_*dir*_ = 0.1 and trained the fusion and matching layers for 500 steps with a learning rate of 1e-4 to achieve integration and alignment. For all other methods, we followed the official protocols and usage instructions provided by the respective authors:

- MISO: https://github.com/kpcoleman/miso/blob/main/tutorial/tutorial.ipynb
- MultiGATE: https://multigate.readthedocs.io/en/latest/index.html
- SpatialGlue: https://spatialglue-tutorials.readthedocs.io/en/latest/index.html
- SIMO: https://github.com/ZJUFanLab/SIMO/tree/main

For fair comparison, parameters for each method were tuned within the ranges recommended in the original studies. For MISO, pixel_size_raw was estimated using the 50 µm grid resolution and the pixel distance between adjacent spots to align image pixel and physical scales for spot-level image feature extraction. For MultiGATE, the learning rate was set to 1e-3 to ensure stable integration, and model performance showed minimal sensitivity to bp_width (Supplementary Fig. 12), which was therefore set to 400. As SpatialGlue uses data-type-specific weight_factors schemes, we applied the Spatial-epigenome-transcriptome training mode following the official documentation. For SIMO, we followed the original protocol in which four adjacent pixels were merged to construct the spatial transcriptomics data, while the original data were treated as single-cell inputs for mapping. Original clustering labels were incorporated as structural priors during training. We observed that increasing the number of mapped cells led to progressive loss of spatial coherence and disruption of tissue organization (Supplementary Fig. 11), possibly because SIMO is based on a single-cell-to-spatial mapping framework rather than direct spot-level co-embedding integration. Therefore, top_num was set to 3 for result presentation and comparison.

#### Marker–Region Concordance

To assess the agreement between identified cortical plate (CP) clusters and CP-associated molecular patterns, we derived a CP marker enrichment score from ATAC data by aggregating normalized chromatin accessibility signals of the canonical CP markers Tbr1 and Satb2 via the score_genes() function in Scanpy. Marker-enriched spots were defined using quantile-based thresholds. For each threshold, cluster purity was defined as the fraction of cluster spots classified as marker-enriched, while coverage was defined as the fraction of marker-enriched spots captured by the cluster.

#### Efficiency Evaluation

We evaluated computational efficiency on a single NVIDIA GeForce RTX 4090 GPU with 24 GB memory. To minimize the influence of hardware-related variability, the training of each method was repeated five times, and we report the mean and standard deviation to provide a more robust estimate of practical runtime. For methods consisting of multiple computational stages, we recorded stage-specific runtimes and present stacked runtimes.

### Spatial alignment and integration of weakly correlated modalities across resolutions

#### Data description

The multi-omics mouse cerebellum dataset[15] comprises three consecutive tissue slices, each representing a distinct spatial omics modality: MAGIC-seq for spatial transcriptomics, PLATO for spatial proteomics, and MALDI-MSI for spatial metabolomics. The MAGIC-seq slice captured 1677 spatial spots at a resolution of 32 μm, profiling 16,116 genes. The PLATO slice provides high-throughput proteomic profiling aligned with the transcriptomic slice, identifying 5722 protein groups after AI enhancement and quality control. The MALDI-MSI slice achieves higher spatial resolution, capturing 3908 spots with 491 metabolite peaks. Spatial domain annotations are available for all three slices, though slight differences exist in domain distributions across modalities. Notably, the slices with different resolutions exhibit significant discrepancies in spatial coordinates.

#### Hyperparameter settings

In the experiments on the multi-omics mouse cerebellum dataset, we set K=4 for KNN graph construction and employed NDT-based coordinate pre-matching prior to training. The registered coordinates were then used in the matching layer for geometry-constrained alignment. During the pretraining stage, we extracted the top 3000 spatially variable features from both the MAGIC-seq transcriptomic slice and the PLATO proteomic slice. These features were normalized and log-transformed before being input into separate two-layer MCGATE models with hidden dimensions of 512 and 32. The number of attention heads was set to 4, and the models were trained for 500 steps using the Adam optimizer with learning rates of 3e-3 and 1e-3, respectively. Considering the relatively low throughput of the MALDI-MSI metabolomic data, we used all available features as input and employed a single-layer MCGATE model with a 32-dimensional embedding and four attention heads, trained for 500 steps with a learning rate of 5e-4. In the integration and alignment stage, we set *β*_*rec*_ = 50 and *β*_*dir*_ = 0.1, and trained the fusion and matching layers for 200 steps using the Adam optimizer with a learning rate of 3e-3.

## Data availability

All experimental data of this study have already been published and are accessible within the corresponding articles and public repositories. Specifically, the 10X Visium DLPFC dataset is available in the spatialLIBD package[76] (http://spatial.libd.org/spatialLIBD); the osmFISH mouse somatosensory cortex dataset is available at http://linnarssonlab.org/osmFISH/; the CODEX human bladder cancer dataset is available from the Aquila database[77] (https://aquila.cheunglab.org/); the BaristaSeq mouse visual cortex dataset is available from the SODB database[78] (https://gene.ai.tencent.com/SpatialOmics/); the Stereo-seq axolotl regenerative telencephalon dataset is available from the STOmicsDB database[79] (https://db.cngb.org/stomics/artista/); the spatial ATAC–RNA-seq E13 mouse embryo dataset is available at https://ki.se/en/mbb/oligointernode; and the spatial multi-omics mouse cerebellum dataset is available from the original article.

## Code availability

The SpatialFuser framework was implemented in the “spatialFuser” Python package, which is open-source for the research community and can be accessible at https://github.com/liwz-lab/SpatialFuser.

## Acknowledgements

This work was supported by the grants of National Natural Science Foundation of China (92474107 and 32570798), National Key R&D Program of China (2021YFF1200903), Major Project of Guangzhou National Laboratory of China (GZNL2024A01003), and Guangdong Basic and Applied Basic Research Foundation of China (2022B1515120077).

