## Supplementary figures for "SpatialFuser: a unified framework for integrative analysis of unpaired spatial multi-omics data"

**Supplementary Fig. 1.** Comparison of spatial domains by clustering assignments.
**Supplementary Fig. 2.** Benchmarking results of tissue domain identification in the DLPFC dataset using commonly applied clustering methods.
**Supplementary Fig. 3.** Robustness of SpatialFuser clustering accuracy across hyperparameter settings and random seeds.
**Supplementary Fig. 4.** Comparison of tissue domain identification ARI before and after alignment BaristaSeq dataset using SpatialFuser, SPACEL, STAligner, and SALT.
**Supplementary Fig. 5.** UMAP visualizations of BaristaSeq dataset before and after alignment using SpatialFuser, SPACEL, STAligner, and SALT.
**Supplementary Fig. 6.** Hyperparameter search for SPACEL.
**Supplementary Fig. 7.** Hyperparameter search for STAligner.
**Supplementary Fig. 8.** Hyperparameter search for SLAT.
**Supplementary Fig. 9.** Original annotated dEGCs field and mutual top-K neighbors distributions in the Stereo-seq axolotl regenerative telencephalon dataset generated by the Sinkhorn-based matching layer during training.
**Supplementary Fig. 10.** UMAP visualizations of integration results by SpatialFuser and MultiGATE on E13 mouse embryo spatial ATAC–RNA-seq data.
**Supplementary Fig. 11.** Cell type distribution of scRNA-seq and scATAC-seq cells generated from the E13 mouse embryo spatial ATAC–RNA-seq dataset.
**Supplementary Fig. 12.** Integration performance of MultiGATE on the E13 mouse embryo spatial ATAC–RNA-seq dataset.
**Supplementary Fig. 13.** Similarity score distributions from random matching and SpatialFuser matching in pairwise alignments across spatial transcriptomics (MAGIC-seq), proteomics (PLATO), and metabolomics (MALDI-MSI) datasets.
**Supplementary Fig. 14.** Sankey plots showing region type correspondence based on pairwise alignments from SpatialFuser across spatial transcriptomics (MAGIC-seq), proteomics (PLATO), and metabolomics (MALDI-MSI) datasets.

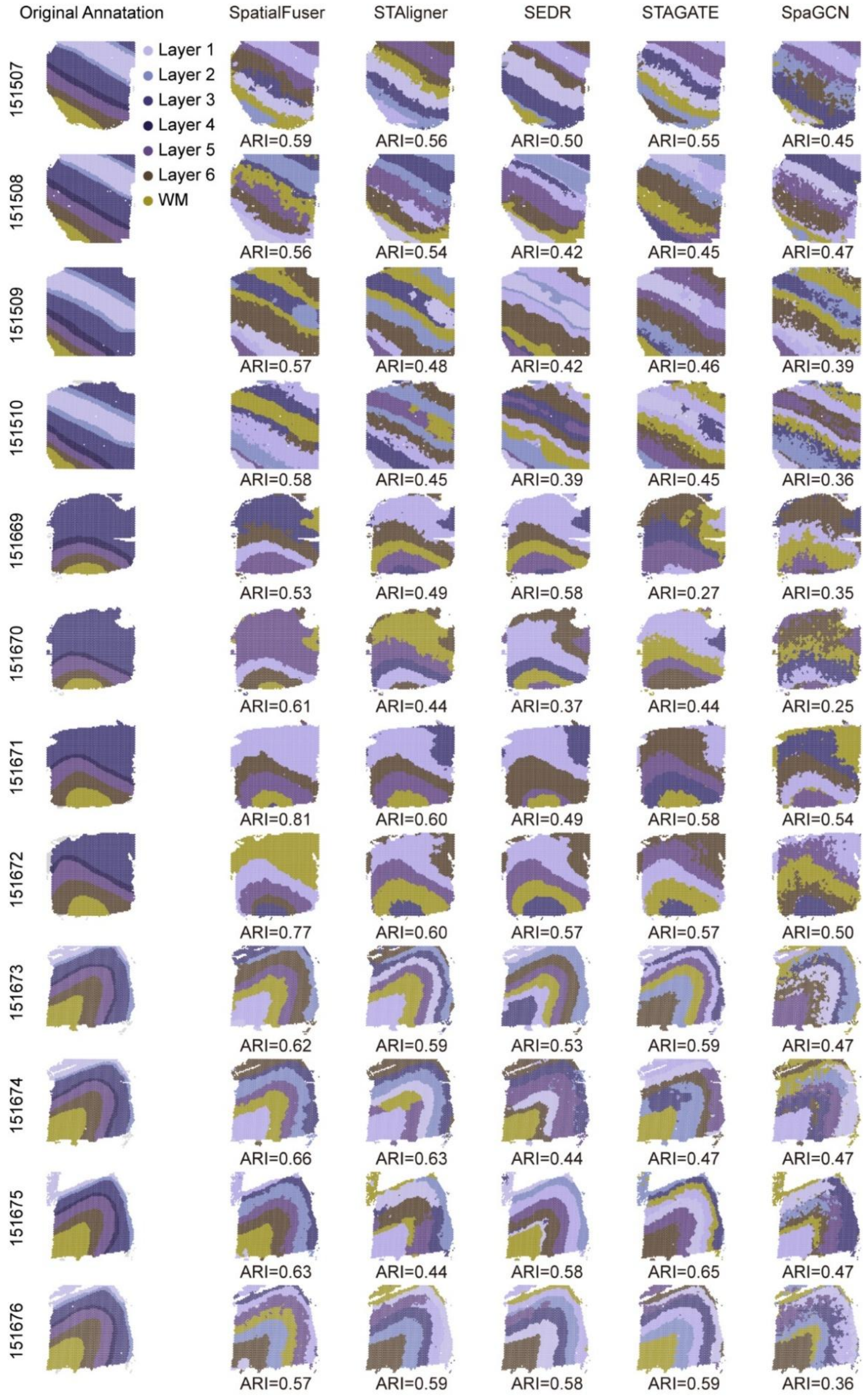


**Supplementary Fig. 1.** Comparison of spatial domains by clustering assignments via SpatialFuser, STAligner, SEDR, STAGATE, SpaGCN and manual annotation in all 12 sections of the DLPFC dataset.

a

b

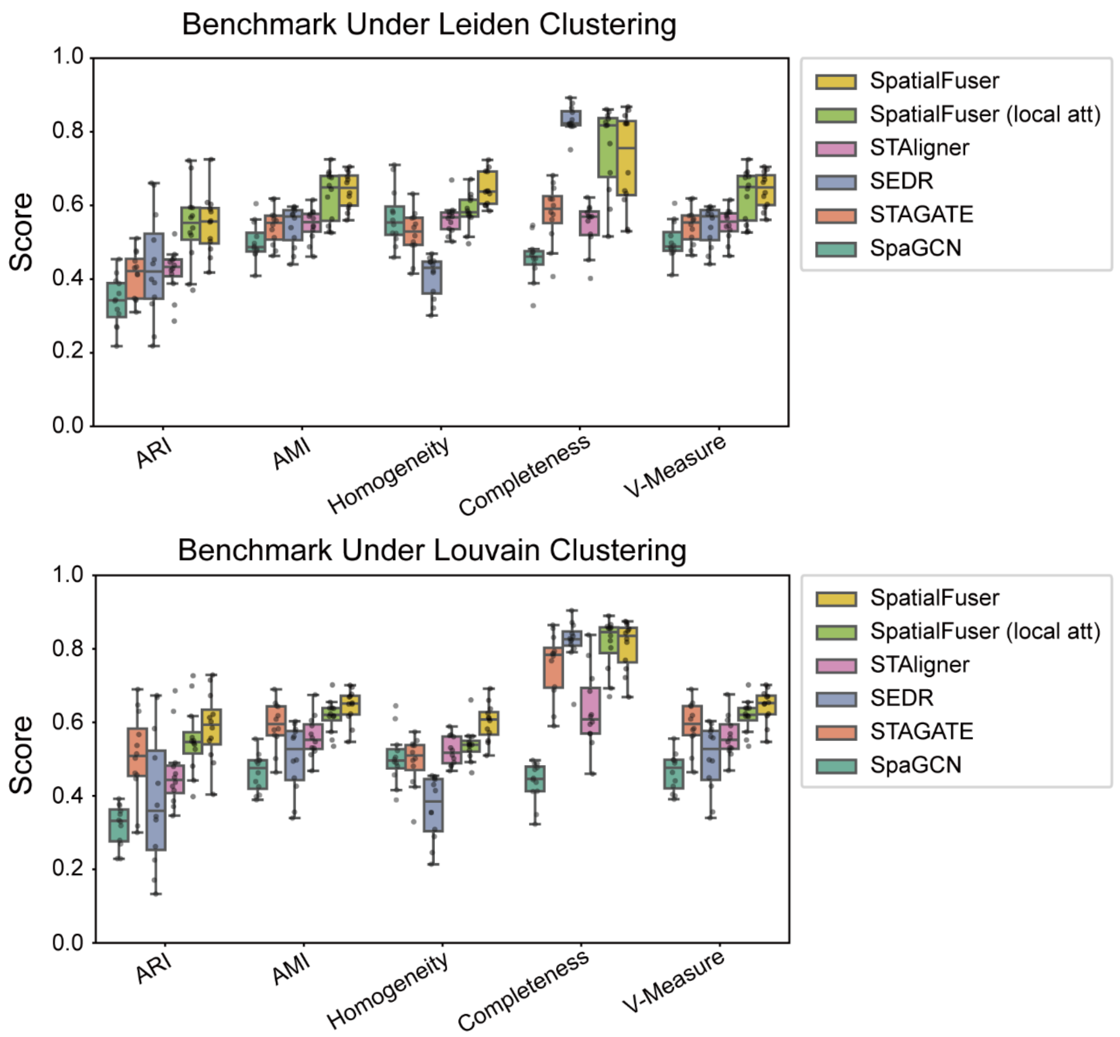


**Supplementary Fig. 2.** Benchmarking results of tissue domain identification in the DLPFC dataset using commonly applied clustering methods. **a,** Boxplots of five evaluation metrics (ARI, AMI, Homogeneity, Completeness, V-measure) for Leiden clustering results across 12 DLPFC sections. **b,** Boxplots of five evaluation metrics (ARI, AMI, Homogeneity, Completeness, V-measure) for Louvain clustering results across 12 DLPFC sections.

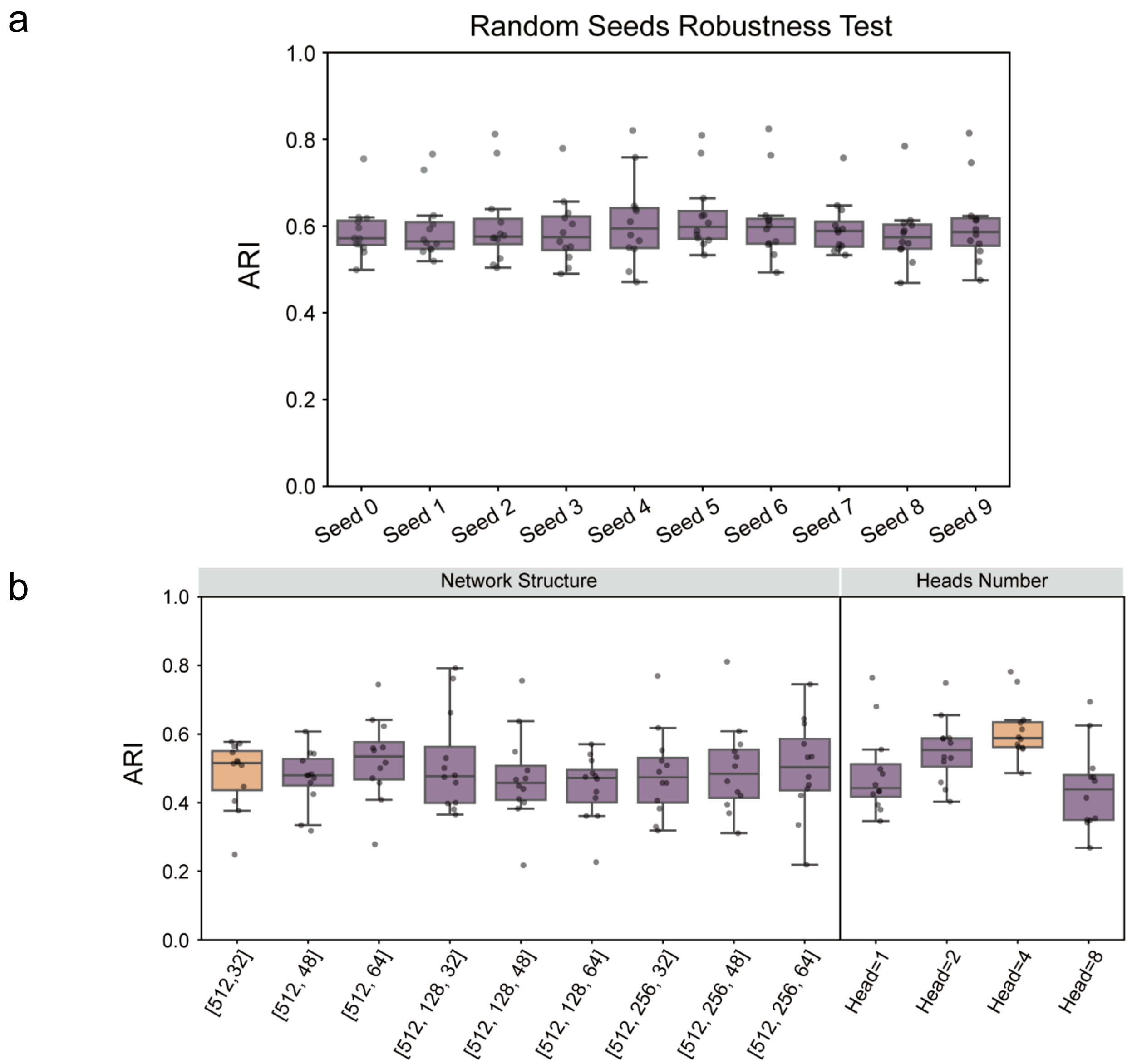


**Supplementary Fig. 3.** Robustness of SpatialFuser clustering accuracy across hyperparameter settings and random seeds. **a,** Clustering ARI of SpatialFuser across all 12 DLPFC sections under different hyperparameter settings. Hyperparameters were tuned via grid search over encoder/decoder layer numbers (2 or 3), hidden layer dimensions (512, 256, or 128), and latent dimensions (32, 48, or 64). **b,** Clustering ARI of SpatialFuser across all 12 DLPFC sections using the default hyperparameters under different random seeds.

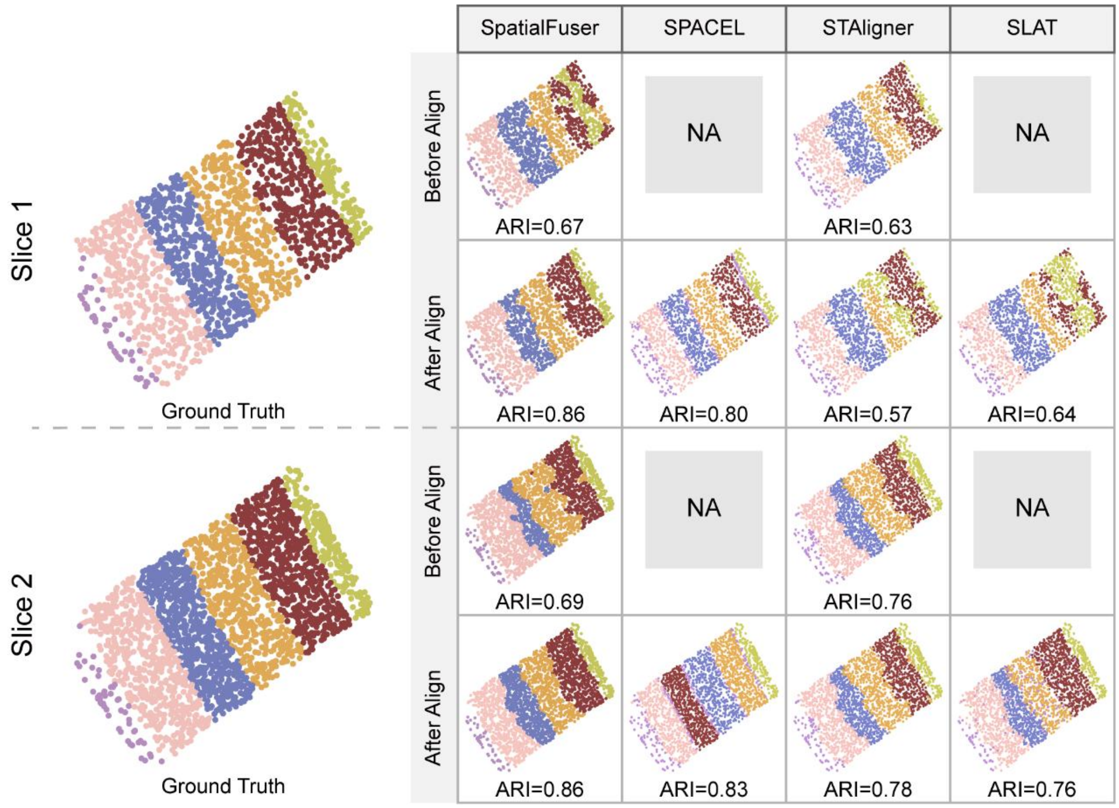


**Supplementary Fig. 4.** Comparison of tissue domain identification ARI before and after alignment in slice 1 and slice 2 of BaristaSeq dataset using SpatialFuser, SPACEL, STAligner, and SALT.

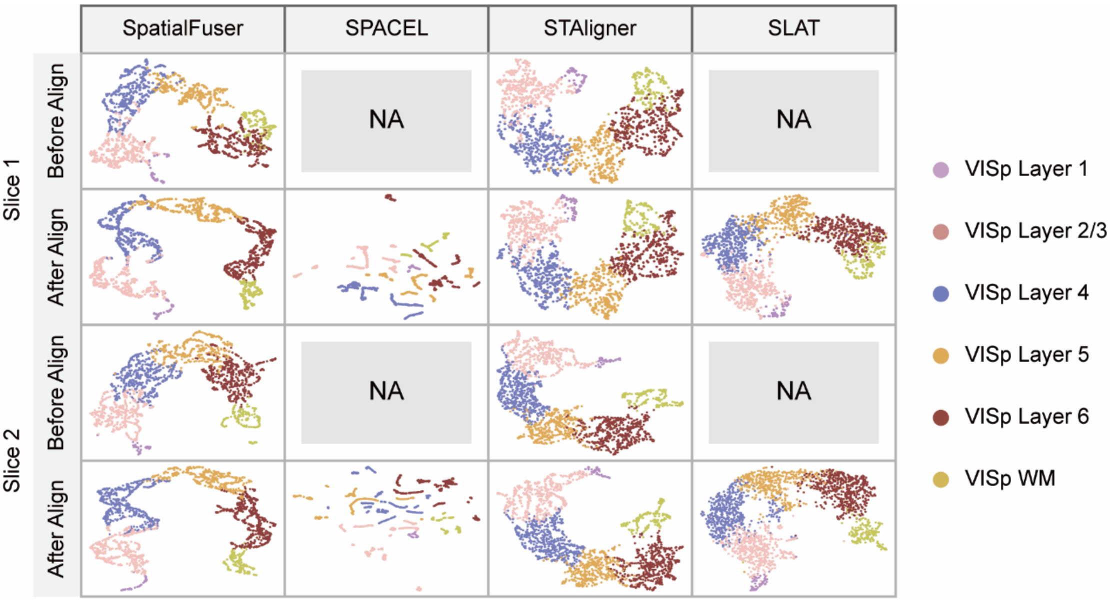


**Supplementary Fig. 5.** UMAP visualizations of slice 1 and slice 2 of BaristaSeq dataset before and after alignment using SpatialFuser, SPACEL, STAligner, and SALT. Spots are colored according to the manual annotations.

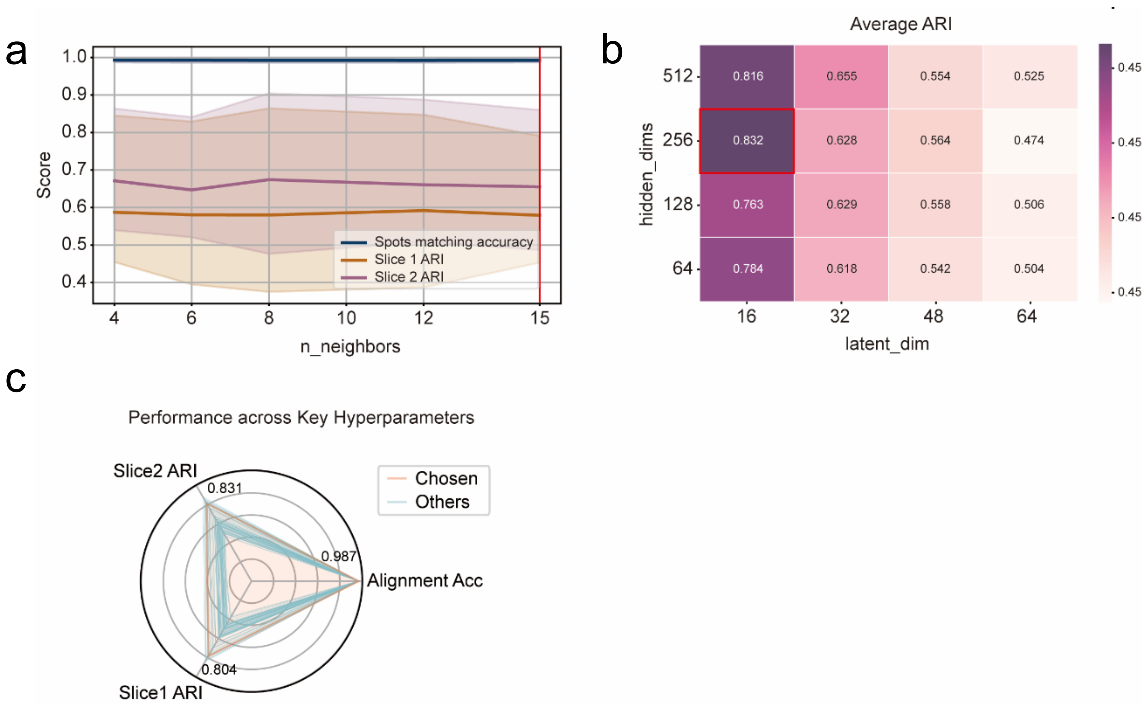


**Supplementary Fig. 6.** Hyperparameter search for SPACEL. **a,** Tissue domain detection performance (ARI) and spot type matching accuracy under different values of n_neighbors. **b,** Average tissue domain detection performance (ARI) across different network structures (hidden_dims and latent_dim). **c,** Radar plots of performance (single-slice domain detection ARI and cross-slice spot matching accuracy) across all grid search parameter combinations. The red line indicates the optimal parameter setting selected.

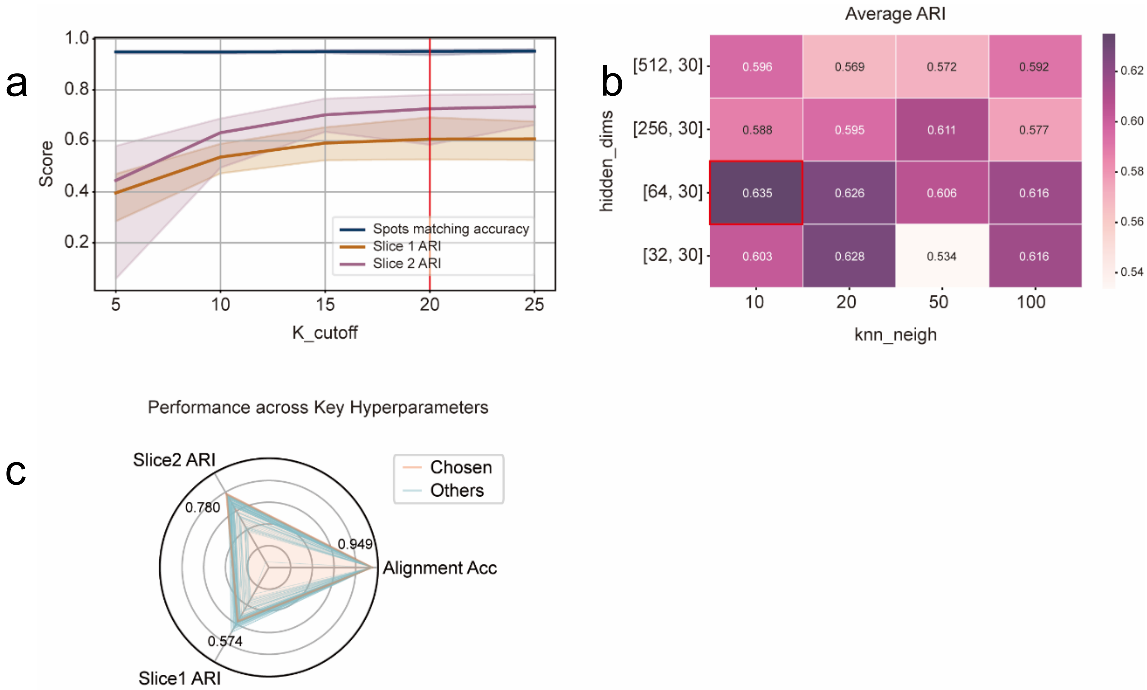


**Supplementary Fig. 7.** Hyperparameter search for STAligner. **a,** Tissue domain detection performance (ARI) and spot type matching accuracy under different values of K_cutoff. **b,** Average tissue domain detection performance (ARI) across different network setting (hidden_dims and knn_neigh). **c,** Radar plots of performance (single-slice domain detection ARI and cross-slice spot matching accuracy) across all grid search parameter combinations. The red line indicates the optimal parameter setting selected.

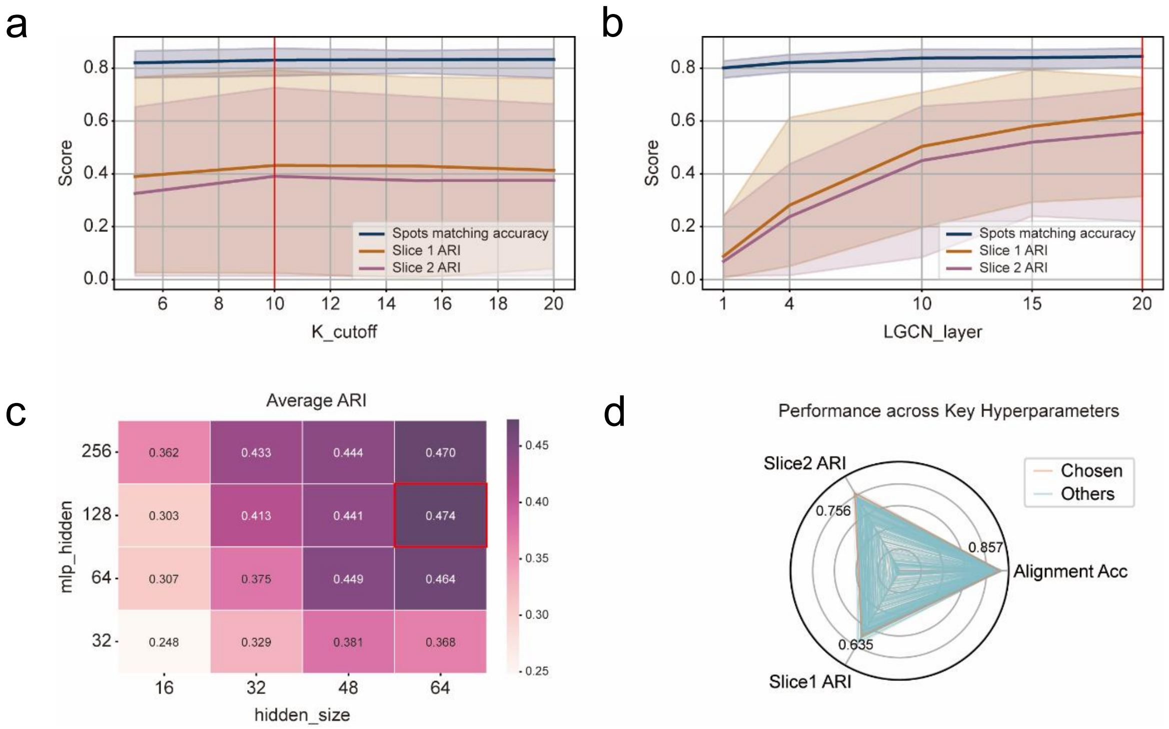


**Supplementary Fig. 8.** Hyperparameter search for SLAT. **a,** Tissue domain detection performance (ARI) and spot type matching accuracy under different values of K_cutoff. **b,** Tissue domain detection performance (ARI) and spot type matching accuracy under different numbers of LGCN_layers (1 and 4, as suggested by the authors). **c,** Average tissue domain detection performance (ARI) across different network structures (mlp_hidden and hidden_size). **d,** Radar plots of performance (single-slice domain detection ARI and cross-slice spot matching accuracy) across all grid search parameter combinations. The red line indicates the optimal parameter setting selected.

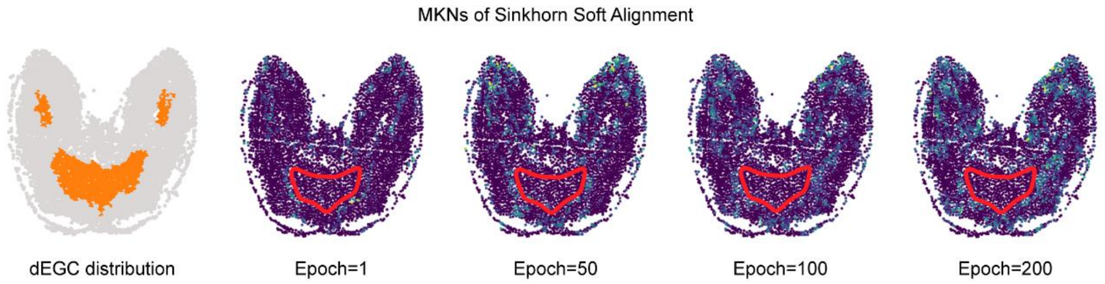


**Supplementary Fig. 9.** Original annotated dEGCs field and mutual top-K neighbors distributions in the Stage 57 slice of the Stereo-seq axolotl regenerative telencephalon dataset generated by the Sinkhorn-based matching layer during training. The green box highlights the dEGCs region, which consistently lacks matched points across multiple training steps (steps 1, 50, 100, and 200).

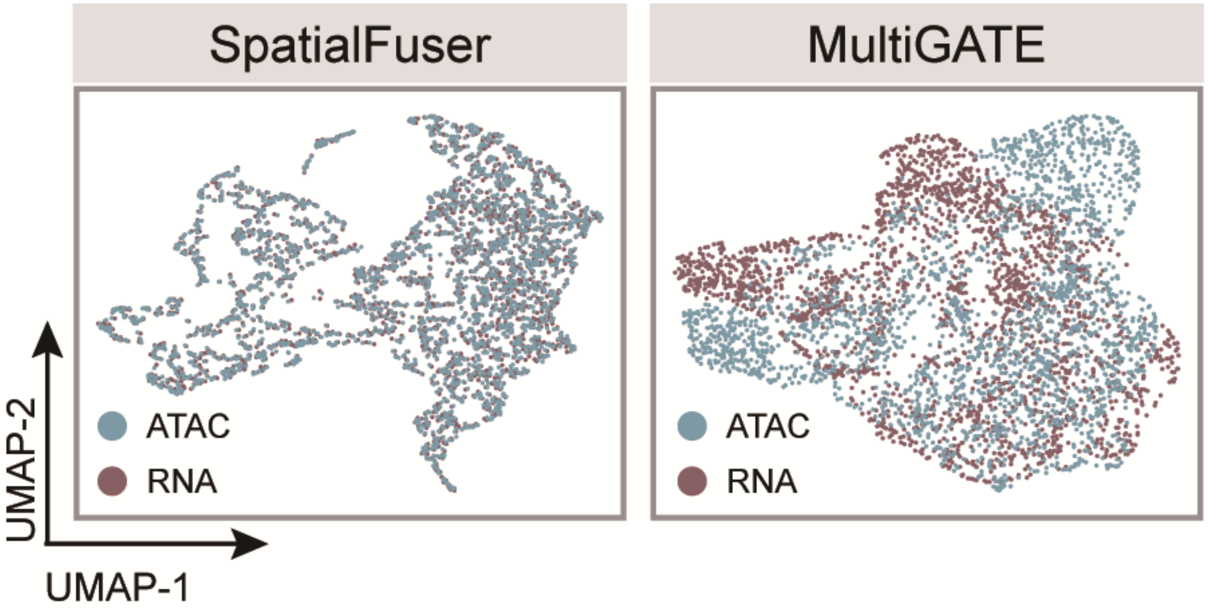


**Supplementary Fig. 10.** UMAP visualizations of integration results by SpatialFuser and MultiGATE on E13 mouse embryo spatial ATAC–RNA-seq data, coloured by modality labels.

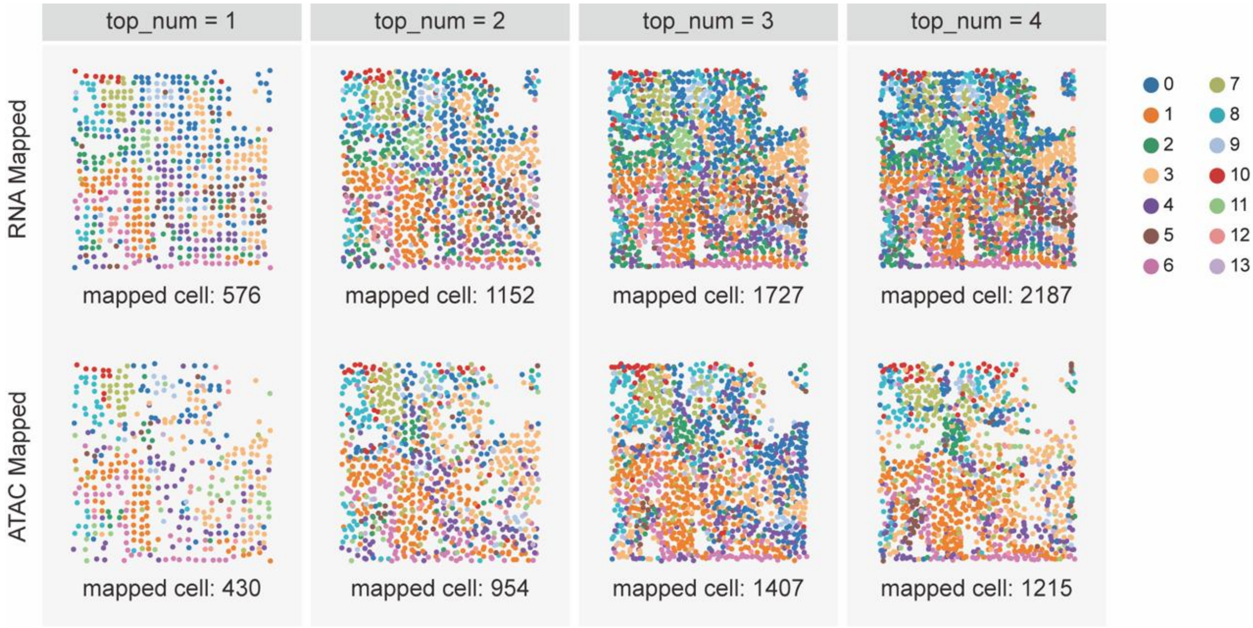


**Supplementary Fig. 11.** Cell type distribution of scRNA-seq and scATAC-seq cells generated from the E13 mouse embryo spatial ATAC–RNA-seq dataset, mapped by SIMO under different values of top_num, and coloured by the original joint cluster labels.

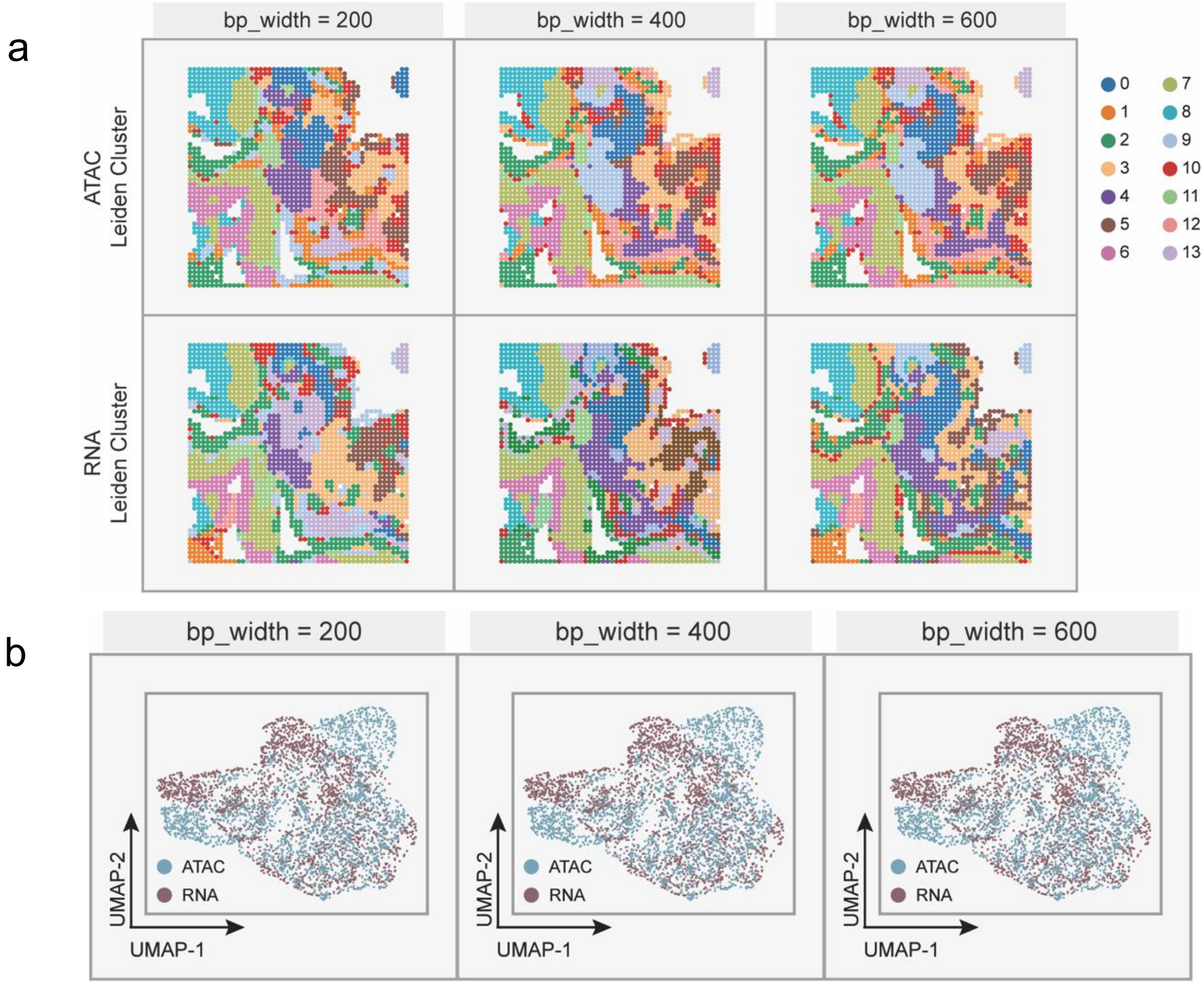


**Supplementary Fig. 12.** Integration performance of MultiGATE on the E13 mouse embryo spatial ATAC–RNA-seq dataset across different bp_width setting (200, 400, 600). **a,** Spatial domains identified by Leiden clustering on low-dimensional embeddings of ATAC (top) and RNA (bottom) generated by MultiGATE under different bp_width settings. **b,** UMAP visualizations of MultiGATE integration results across different bp_width values, colored by modality labels.

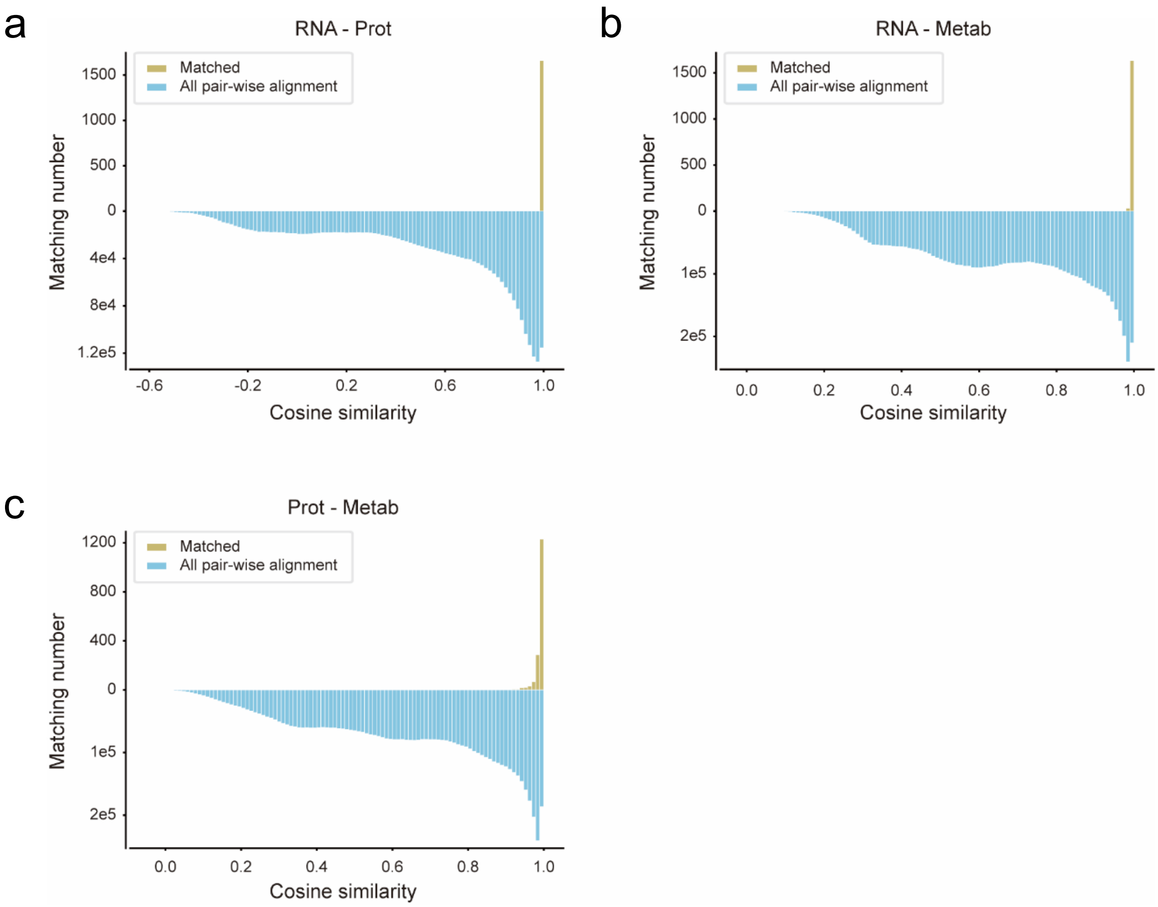


**Supplementary Fig. 13.** Similarity score distributions from random matching and SpatialFuser matching in pairwise alignments across spatial transcriptomics (MAGIC-seq), proteomics (PLATO), and metabolomics (MALDI-MSI) datasets. **a,** Similarity score distribution in MAGIC-seq and PLATO alignment. **b,** Similarity score distribution in MAGIC-seq and MALDI-MSI alignment. **c,** Similarity score distribution in PLATO and MALDI-MSI alignment.

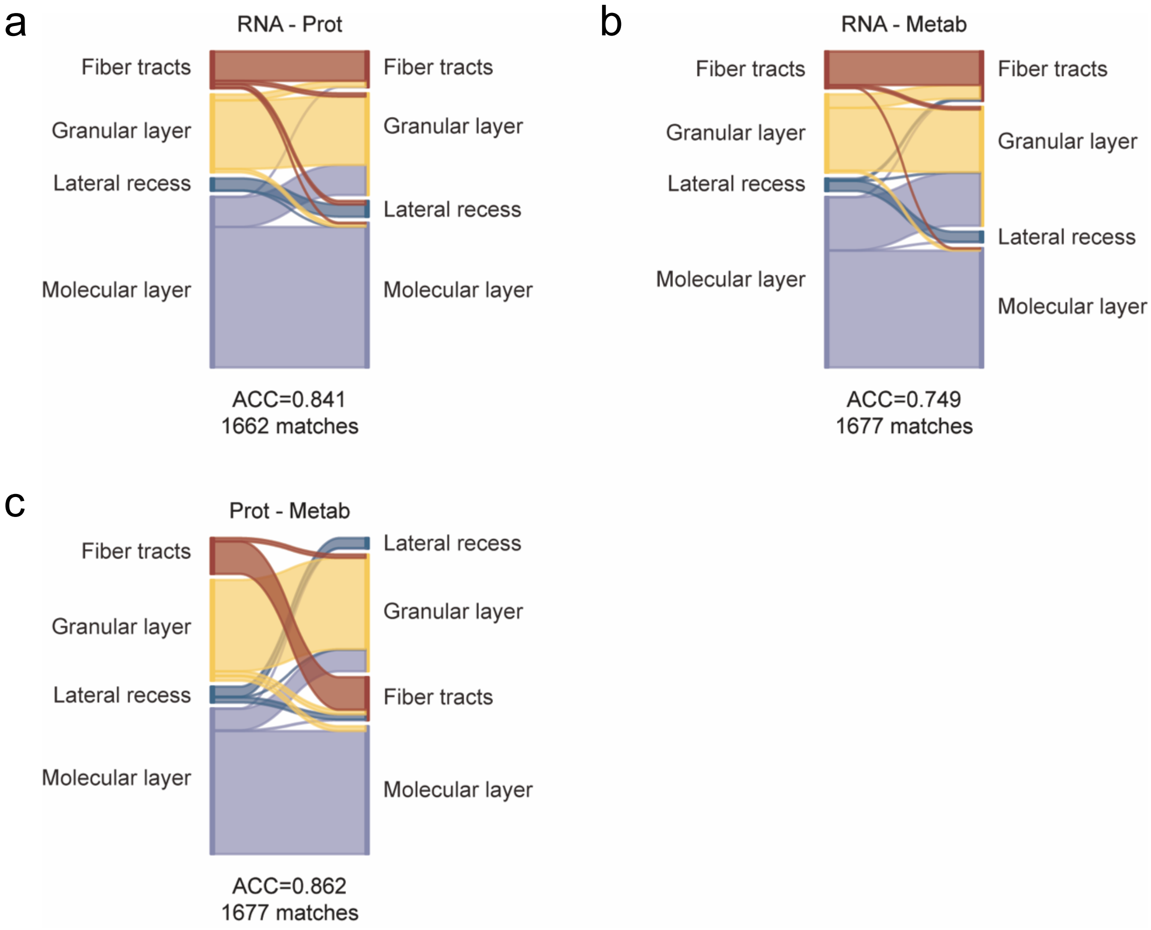


**Supplementary Fig. 14.** Sankey plots showing region type correspondence based on pairwise alignments from SpatialFuser across spatial transcriptomics (MAGIC-seq), proteomics (PLATO), and metabolomics (MALDI-MSI) datasets. **a,** Spot type matching in MAGIC-seq and PLATO alignment. **b,** Spot type matching in MAGIC-seq and MALDI-MSI alignment. **c,** Spot type matching in PLATO and MALDI-MSI alignment.
